# RamEx: An R package for high-throughput microbial ramanome analyses with accurate quality assessment

**DOI:** 10.1101/2025.03.10.642505

**Authors:** Yanmei Zhang, Gongchao Jing, Rongze Chen, Yanhai Gong, Yuandong Li, Yongshun Wang, Xixian Wang, Jia Zhang, Yuli Mao, Yuehui He, Xiaoshan Zheng, Mingchao Wang, Hao Yuan, Jian Xu, Luyang Sun

## Abstract

Microbial single-cell Raman spectroscopy (SCRS) has emerged as a powerful tool for label-free phenotyping, enabling rapid characterization of microbial diversity, metabolic states, and functional interactions within complex communities. However, high-throughput SCRS datasets often contain spectral anomalies from noise and fluorescence interference, which obscure microbial signatures and hinder accurate classification. Robust algorithms for outlier detection and microbial ramanome analysis remain underdeveloped. Here, we introduce RamEx, an R package specifically designed for high-throughput microbial ramanome analyses with robust quality control and phenotypic classification. At the core of RamEx is the Iterative Convolutional Outlier Detection (ICOD) algorithm, which dynamically detects spectral anomalies without requiring predefined thresholds. Benchmarking on both simulated and real microbial datasets—including pathogenic bacteria, probiotic strains, and yeast fermentation populations—demonstrated that ICOD achieves an F1 score of 0.97 on simulated datasets and 0.74 on real datasets, outperforming existing approaches by at least 19.8%. Beyond anomaly detection, RamEx provides a modular and scalable workflow for microbial phenotype differentiation, taxonomic marker identification, metabolic fingerprinting, and intra-population heterogeneity analysis. It integrates Raman-based species-specific biomarkers, enabling precise classification of microbial communities and facilitating functional trait mapping at the single-cell level. To support large-scale studies, RamEx incorporates C++ acceleration, GPU parallelization, and optimized memory management, enabling the rapid processing of over one million microbial spectra within an hour. By bridging the gap between high-throughput Raman-based microbial phenotyping and computational analysis, RamEx provides a comprehensive toolkit for exploring microbial ecology, metabolic interactions, and antibiotic susceptibility at the single-cell resolution. RamEx is freely available under the MIT license at https://github.com/qibebt-bioinfo/RamEx.

## Introduction

Single-cell Raman spectroscopy (SCRS) has emerged as a powerful tool for microbial metabolic phenotyping, as enabling label-free, non-invasive characterization of microbial cells based on their unique vibrational “fingerprints” [1–3]. By capturing metabolic states at the single-cell level, SCRS has become an essential component of the expanding microbial single-cell “-omics” landscape [4, 5], enabling direct investigation of complex microbial communities, ranging from microbial identification [6–8] and biomass quantification [9–11] to environmental stress evaluation [12, 13] and metabolic heterogeneity analysis[14, 15]. With single-cell research generating unprecedented volumes of data, recent technological advances in high sensitivity detectors [16, 17] and microfluid-based cell sorting Raman measurement systems [18–22] have significantly enhanced the throughput and scalability of SCRS. Notably, modern Raman platforms now enable high-throughput microbial phenotyping, capturing thousands of Raman spectra per minute [23, 24], enabling the rapid acquisition of large-scale microbial Raman datasets [25–27]. Concurrently, the development of standardized Raman spectral databases [28] has facilitated comparative microbial analyses and reproducibility across studies. These studies highlight the potential of Raman-activated cell sorting for identifying and screening key players in targeted processes.[29, 30] However, despite these technological strides, computational tools to extract meaningful insights from the deluge of microbial Raman data have remained underdeveloped, presenting critical bottlenecks to the field.

Several software packages and algorithms have been developed to support microbial Raman spectral data analysis, including both commercial platforms and open-source tools (**Supplementary table 1**). Commercial platforms such as LabSpec, WiRE, and WITec are user-friendly and allow non-specialists to perform basic preprocessing and spectral feature identification. However, they are largely confined to elementary preprocessing tasks and lack the scalability and advanced data mining capabilities required for large-scale, high-dimensional biological applications [31, 32]. Open-source packages, such as MCR-ALS [33], MALDIquant [34], and RamanSpy [35], provide greater flexibility and support more complex workflows, particularly in leveraging community-driven resources and facilitating seamless integration with modern framework. In addition, deep-neuron network has been used to perform the preprocessing steps for Raman spectrum and observed considerable results with traditional methods [36–38]. However, existing tools are not specifically optimized for microbial ramanome analysis, particularly in addressing key challenges such as microbial population clustering, phenotypic heterogeneity, and metabolic interaction networks[39]. An effective approach for handling large-scale ramanome data is to adapt methodologies from other omics disciplines, particularly single-cell sequencing. For instance, scRNA-seq analysis tools such as Seurat [40], BayesSpace [41], and Monocle [42] have been applied for tissue structure reconstruction [43], and chemometric spectral unmixing techniques have been repurposed for compositional analysis [33]. However, these adapted approaches often fail to capture the unique characteristics of microbial Raman spectra, such as continuous spectral signals with baseline variations, distinctive noise patterns, and overlapping signals [44, 45]. Unlike single-cell transcriptomics, which operates on sparse, zero-inflated sequencing data, microbial ramanome data follows a fundamentally different statistical distribution, making direct adaptation of sequencing-based approaches suboptimal for microbial spectral analysis [46, 47].

Furthermore, due to the inherent detection characteristics of Raman spectroscopy, various spurious signals such as thermal noise, shot noise, black-body radiation from samples, fluorescence from cellular structural features, cell motion artifacts, and environmental fluctuations can easily distort the weak signals originating from biological tissues [48–50]. Although the optimized design of optical system elements can reduce the frequency of these irrelevant signals to some extent, such interference remains unavoidable during automated high-throughput data acquisition [51, 52]. Improper removal of these anomalous spectra can severely compromise downstream phenotype analyses, particularly in complex microbial communities[39, 53, 54]. However, few outlier detection algorithms specifically designed for large-scale microbial ramanomes. Most existing methods [55–57] rely on empirically determined thresholds that require manual optimization for different experimental conditions, making them poorly suited for the complex, heterogeneous nature of microbial spectral data [58]. While machine learning approaches [50, 59], particularly deep learning methods, have demonstrated promising results in spectral analysis but necessitate extensive training datasets and careful hyperparameter optimization [6, 60], limiting their generalizability across different microbial communities, experimental platforms, and environmental conditions.

To address these pressing challenges in high-throughput microbial Raman spectral analysis, we present Ramanome Explorer (RamEx), an integrated computational framework specifically designed for large-scale microbial ramanomes processing and analysis. At its core, RamEx introduces novel algorithmic advancements for robust spectral quality control, microbial phenotype differentiation, and metabolic fingerprinting, optimizing data extraction from high-dimensional microbial Raman spectra. Its scalable architecture ensures efficient analysis of large and diverse microbial datasets, as demonstrated in benchmarking tests across eight biological datasets (cumulative >1 million spectra) spanning multiple microbial taxa and environmental conditions (**Supplementary table 2**). RamEx provides a standardized yet flexible analytical framework, combining automated data processing pipelines with customizable workflow modules to accommodate a broad range of microbial applications.

By bridging the current technical and infrastructural gaps in microbial Raman-omics, RamEx represents a crucial step towards establishing Raman flow cytometry as a mainstream technology, and opens new possibilities for large-scale functional microbial profiling at unprecedented scales.

## Results

### RamEx provides a versatile multifaceted framework for microbial ramanome analysis

Understanding complex and high-dimensional single-cell Raman spectroscopy datasets presents significant challenges due to issues like high collinearity, nonlinearity, and the presence of outliers. To address the challenges in analyzing and interpreting ramanome data, we developed RamEx, an R package designed specifically to analyze large-scale Raman datasets using novel algorithms and comprehensive single-cell analysis workflows. It features: (*i*) an outlier detection algorithm that operates without prior knowledge or fixed criteria; (*ii*) optimized clustering and marker identification algorithms adapted to the unique properties of high dimensional Raman spectra; (*iii*) curated computational framework with tools and pipelines for key Raman tasks such as cell type/species identification, clustering phenotypic analysis, antibiotic resistance detection and molecular composition analysis; (*iv*) enhanced computing efficiency through C++ optimization and GPU computing; and (*v*) standardized Raman dataset format with integrated metadata and evaluation metrics. RamEx is freely available at GitHub (https://github.com/qibebt-bioinfo/RamEx).

RamEx is structured into three core modules: the basic module, the spectral preprocessing module, and the data analysis and modeling module (**Fig. 1**), encompassing a variety of useful analytical functionalities in metabolic phenotyping of cells (**Supplementary table 3**). First of all, the basic module handles data import, data evaluation, and data management, supporting cross-platform input formats from mainstream instrument manufacturers such as Horiba, Renishaw, Thermo Fisher Scientific, WITec, and Bruker. It efficiently manages both single-point data collection, where each spectrum is stored in a separate file, and mapping data enriched with coordinated information. The module automates data type organization, eliminating the need for user intervention or configuration. It also introduces the Raman Attribute Table, which assesses datasets based on reproducibility, interpretability, Raman entropy, signal-to-noise ratio (SNR), and diversity. This tool enables users to efficiently evaluate the quality and potential of large datasets for subsequent analysis (see **Methods**; **Supplementary Fig. 1**).

**Figure 1.**
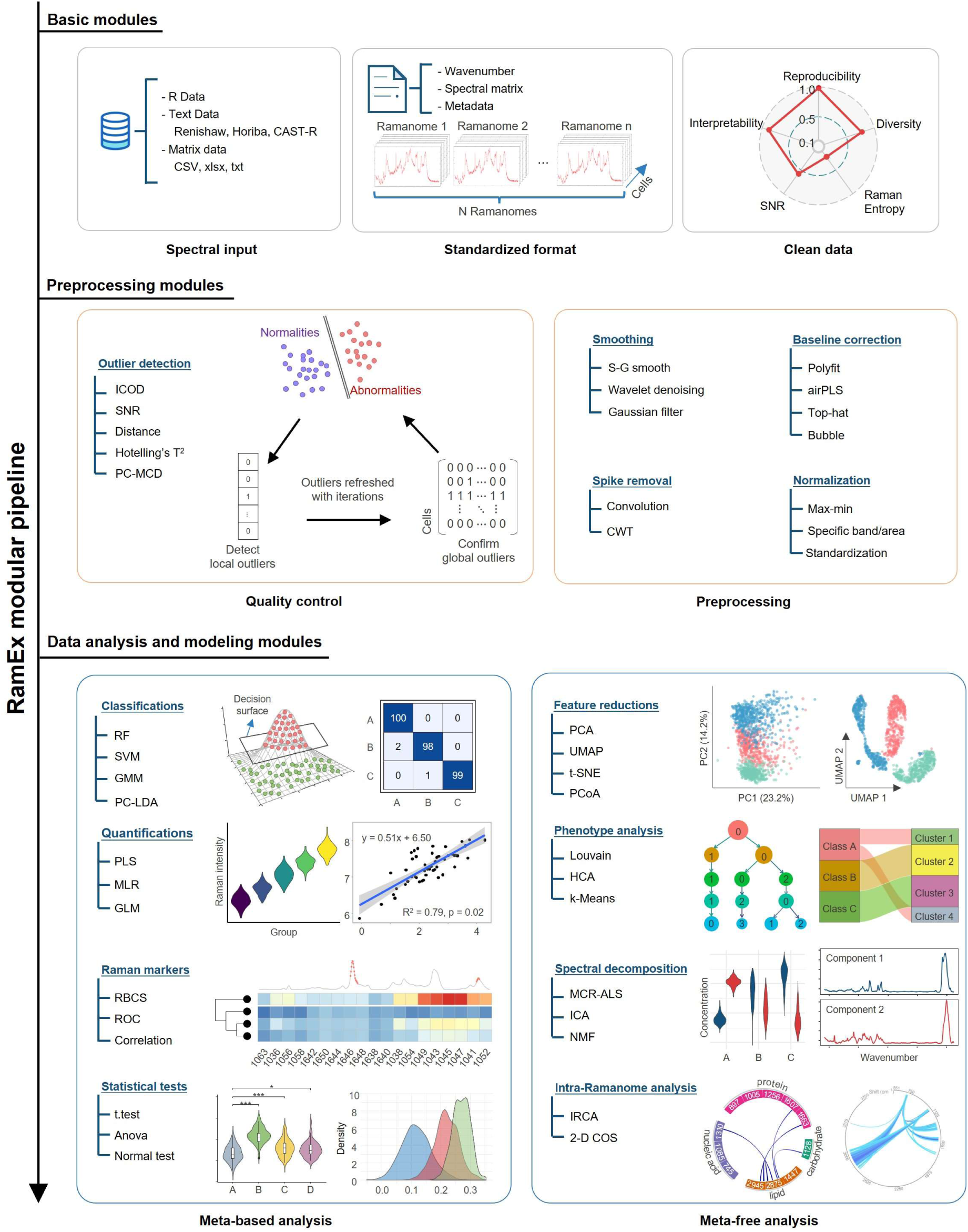
Schematic overview of the RamEx computational framework. The framework comprises three core modules: basic module (input/output management and data format standardization and evaluation), preprocessing modules (quality control and preprocessing steps), and data analysis and modeling modules (classification, quantification, marker identification, phenotype analysis, spectral decomposition and intra-ramanome analysis)

Moreover, the preprocessing module includes a novel outlier detection algorithm we developed called Iterative Convolutional Outlier Detection (ICOD), which enhances data quality by effectively identifying aberrant spectra without requiring empirical threshold settings. RamEx also provides methods for detecting other common outliers, offering flexibility and adaptability in quality control (QC). Beyond outlier detection, the module offers a carefully selected range of classic spectral preprocessing tools, including spike removal, smoothing, baseline correction, and normalization. Multiple methods are provided for each step to suit specific dataset characteristics. Users can customize methods and parameters for comprehensive adjustment or utilize a standardized workflow for rapid analysis, ensuring adaptability and efficiency in processing Raman spectra.

Finally, the data analysis and modeling module provides a suite of functions supporting the primary applications of ramanomics, broadly categorized into meta-based (supervised) and meta-free (unsupervised) applications. For meta-based analyses, RamEx offers classification methods such as Principal Component Analysis–Linear Discriminant Analysis (PCA-LDA), Random Forest, Support Vector Machines (SVM), and so on, as well as quantitative analysis tools like Partial Least Squares Regression (PLS), Multiple Linear Regression (MLR), and Generalized Linear Regression (GLR). Marker identification is optimized for both qualitative and quantitative scenarios, employing both machine learning-based recognitions (e.g., Raman Barcode of Cellular-response to Stresses, RBCS [61]) and univariate identifications based on band traversal (e.g., correlation-based or ROC-based approaches).

For meta-free applications, the module includes dimensionality reduction algorithms such as Uniform Manifold Approximation and Projection (UMAP) and t-distributed Stochastic Neighbor Embedding (t-SNE), which are exceptional for visualizing high-dimensional, highly correlated, and subtly varying biological spectra. Unsupervised clustering algorithms provided include Gaussian Mixture Model (GMM), hierarchical clustering, Louvain, and Leiden algorithms, addressing diverse user requirements and enabling comprehensive comparisons of algorithmic results. Unique to spectral analysis, RamEx incorporates spectral decomposition methods such as Multivariate Curve Resolution–Alternating Least Squares (MCR-ALS), Independent Component Analysis (ICA), and Non-negative Matrix Factorization (NMF) for chemical interpretations. Additionally, we incorporated the Intra-Ramanome Correlation Network Analysis (IRCA) algorithm, specifically designed to capture the subtle but biological meaningful metabolic heterogeneity among Raman spectra from individual cells within a microbial population, can detect the interconversions among metabolites [62].

Overall, RamEx offers a comprehensive suite of user-friendly tools, strategically integrated to facilitate the effective analysis and interpretation of complex Raman data. The subsequent section demonstrates the practical application of these methods.

### Robust quality control in RamEx ensures high-fidelity microbial ramanome analysis

Identifying abnormal spectral acquisitions is crucial for enhancing data quality in biological single-cell datasets. Currently, this process often involves using a combination of filtering criteria and adjusting empirical thresholds tailored to specific projects. The challenge of setting these thresholds intensifies when integrating datasets from multiple batches, as batch variations require meticulous calibration to ensure consistent and reliable analysis outcomes. In single-cell Raman spectroscopy, biological Raman spectra generally display consistent dimensions (number of bands) and similar patterns across different groups, resulting in a lack of obvious variation. This similarity complicates automated quality assessment, such as spectral similarities and quality parameters, which rely on pre-defined thresholds. Moreover, these criteria are highly empirical and often require expert judgment. To address this, we developed the ICOD algorithm in RamEx, which dynamically identify and remove low-quality Raman spectra iteratively, without relying on predefined criteria. By assuming that most biological spectra within a population share consistent overall features, it iteratively identifies Raman outliers in an unsupervised manner, with unbiased consideration of all meaningful peaks (**Fig. 2a**, see **Methods**).

**Figure 2.**
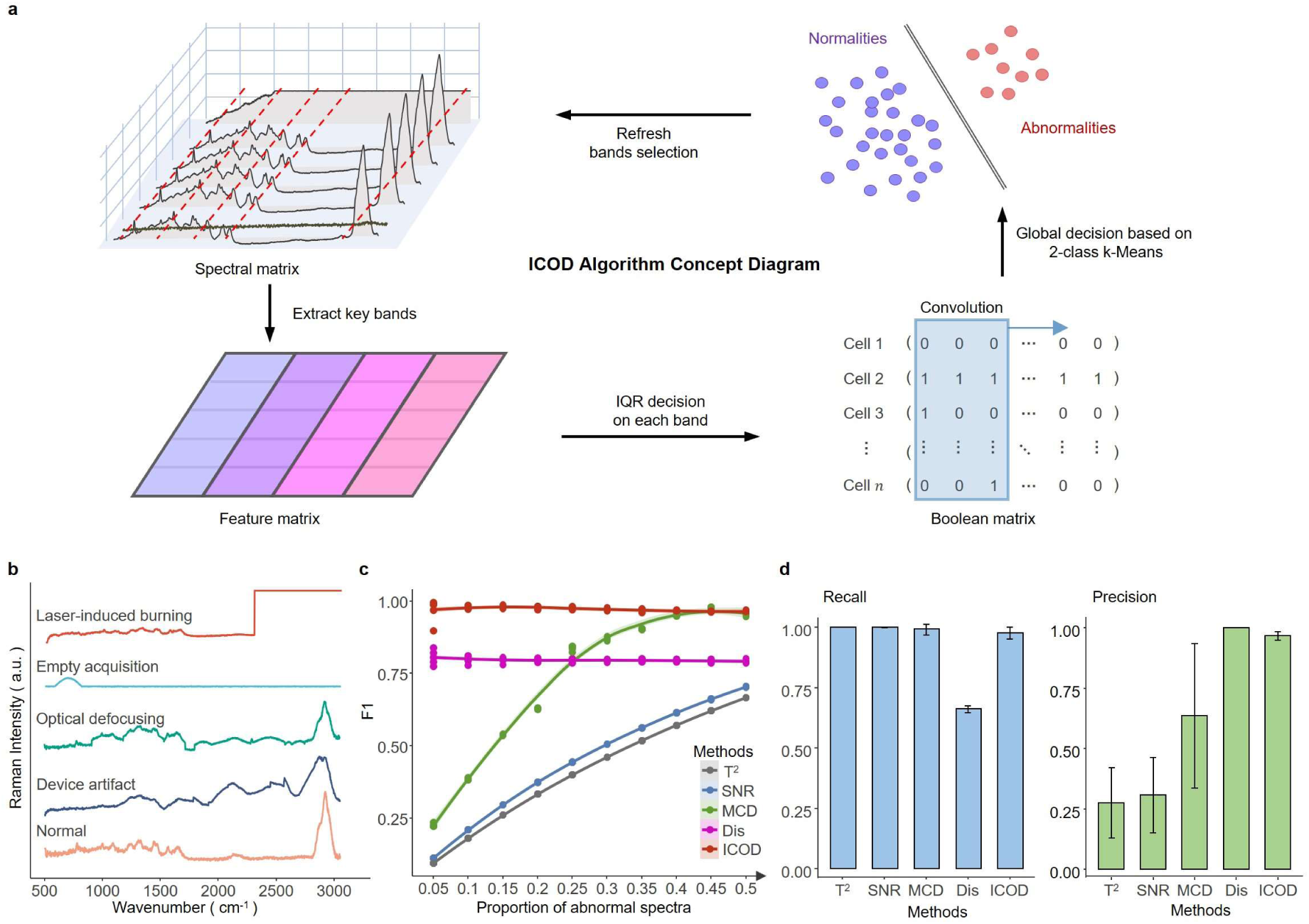
ICOD is robust to various quality levels. **a**, Schematic illustration of ICOD highlighting key computational steps. **b**, Types of simulated anomaly spectra. **c**, F1 scores of five outlier detection methods across datasets with varying proportions of abnormalities. **d**, Recall and precision of different outlier detection methods, where dots represent means; error bars indicate standard deviations, n = 5 independent datasets across 10 quality levels.

Specifically, the ICOD method employs an iterative unsupervised clustering approach to effectively identify and classify outliers in single-cell Raman spectra. Initially, peak identification is conducted using a modified bubble method [63], which isolates intensity information at peak positions, thereby reducing the data’s dimensionality from thousands of features to a few dozen. This reduction not only decreases computational load but also retains critical spectral information while mitigating background noise. The original dataset *X_n_*_×*m*_ will be transformed into 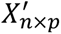, where *p* ≪ *m*. Subsequently, the quartile outlier method is applied to detect anomalies at each peak, where an observation *x_ij_* is considered a local outlier if *x_ij_* < *Q*1*_j_* − 1.5 × *IQR_j_* or *x_ij_* > *Q*3*_j_* + 1.5 × *IQR_j_*. This process converts the dataset into a Boolean matrix *B_n_*_×*p*_, where *B_ij_* = 1 if *x_ij_* is an outlier, and *B_ij_* = 0 otherwise. This matrix is then subjected to k-means clustering and convolution filtering, with *k* = 2, to distinguish spectra into normal and anomalous categories. Convolution enhances the robustness of clustering by diminishing the effects of local anomalies, while a global outlier vector is iteratively refined until convergence. In this global outlier vector, values are dynamically adjusted to reflect the probability of being an outlier, with higher values indicating a greater likelihood. The iterative update process further refines classification by recalculating the matrix of spectra identified as normal in each cycle, thus enhancing outlier detection accuracy through continuous recalibration. This iterative refinement progressively sharpens the distinction between normal and anomalous spectra by accounting for the inherent redundancy in Raman signals, which effectively balances the weights of different peak regions to prevent broader bands (such as CH region at 2750-3000 cm⁻¹) from overshadowing narrower peaks (like the phenylalanine region around 1003 cm^-1^) during outlier detection.

To demonstrate the performance of ICOD, we conducted a comparative analysis against several established outlier detection techniques, including threshold-based approaches like Euclidean distance [55] and signal-to-noise ratio (SNR) [56] (to remove samples with low similarity and low SNR), statistics-based approaches like Hotelling’s T^2^ [59, 64] and Principal components - minimum covariance determinant (PC-MCD) [65] (to remove samples with low confidence), using both simulated large-scale datasets and real benchmarking studies (**Methods**). Four kinds of common anomalous spectra [55] - laser-induced burning, empty acquisition, optical defocusing, and device artifact - along with high-quality Raman spectra, were generated using a Gaussian Mixture Model (**Fig. 2b**, **Supplementary Fig. 2a**). Each test dataset consisted of 10,000 simulated Raman spectra, with the proportion of anomalous spectra varying from 5% to 95%, resulting in 19 distinct datasets. These anomalous spectra were randomly selected from the four afore-mentioned categories. To ensure reproducibility, five replicates were created for each specific percentage of anomalous spectra, culminating in a total of 95 datasets.

Notably, ICOD outperformed other outlier detection methods, consistently achieving a superior F1 score of 0.97 (**Fig. 2c**). In contract, other methods were sensitive to the proportion of outliers, performing poorly at low outlier ratios. Specifically, Hotelling’s T², SNR, and PC-MCD exhibited overly stringent criteria, misclassifying many spectra, while the Euclidean distance method was too permissive. ICOD demonstrated high accuracy and precision due to its iterative convergence process, detecting over 75% of outliers by the second epoch and the rest in subsequent iterations (**Supplementary Fig. 2b**), reflecting ICOD’s adaptive learning rate and robust convergence. Gradient testing revealed ICOD’s enhanced performance at outlier compositions below 70%, applicable to most scenarios (**Fig. 2c** and **Supplementary Fig. 2c**). However, beyond this threshold, anomaly types such as optical defocusing and device artifact predominated, leading to dataset distortion.

To further benchmark ICOD’s performance under real-world conditions, we analyzed several publicly available ramanome datasets, along with a self-collected dataset (**Fig. 3a**). The datasets involved data collected from various Raman instruments, species, and outlier proportions, featuring dry smear samples comprising 9,641 spectra from pathogenic bacteria [13], 49,787 spectra from *Mycobacterium abscessus* under drug exposure, and 6,186 spectra from multiple probiotic strains [66]. Additionally, these datasets included 17,819 spectra from *Escherichia coli* under various treatments [23] and 7,437 spectra related to chlorflavonin production, both obtained using Raman flow cytometry. When compared to expert consensus-determined outlier labels, ICOD consistently outperformed competing algorithms across these datasets in terms of F1 scores, achieving an average score of 0.74, 19.8% higher than the next best method (mean F1 score: 0.61 of SNR method) (**Fig. 3b**). Further analysis shows that ICOD effectively balanced precision and recall in outlier detection across these five datasets (**Fig. 3c**), whereas Hotelling’s T^2^ and PC-MCD were overly stringent, resulting in lower data retention rates, and methods based on distance and SNR lacked sensitivity to anomalies (**Fig. 3b**). These results demonstrated the reliability and adaptability of ICOD as a leading method among those tested. Notably, while the SNR method achieved a better F1 score on the “Bacteria” dataset, a small number of outliers still existed, distorting the overall PCA score space as shown in **Fig. S3a**. From the results of reduction and Adonis analysis, the datasets processed by ICOD exhibited clearer inter-group differences and tighter intra-group clustering (**Supplementary Fig. 3**). Overall, ICOD demonstrated exceptional strengths across both simulated and real-world datasets. However, evaluating their performance solely through F1 scores may obscure its true advantages and does not fully encapsulate the impact of quality control on subsequent data analysis. Next, we further illustrated the benefits of ICOD by applying it to new datasets and showcasing its impact on down-stream analyses within RamEx.

**Figure 3.**
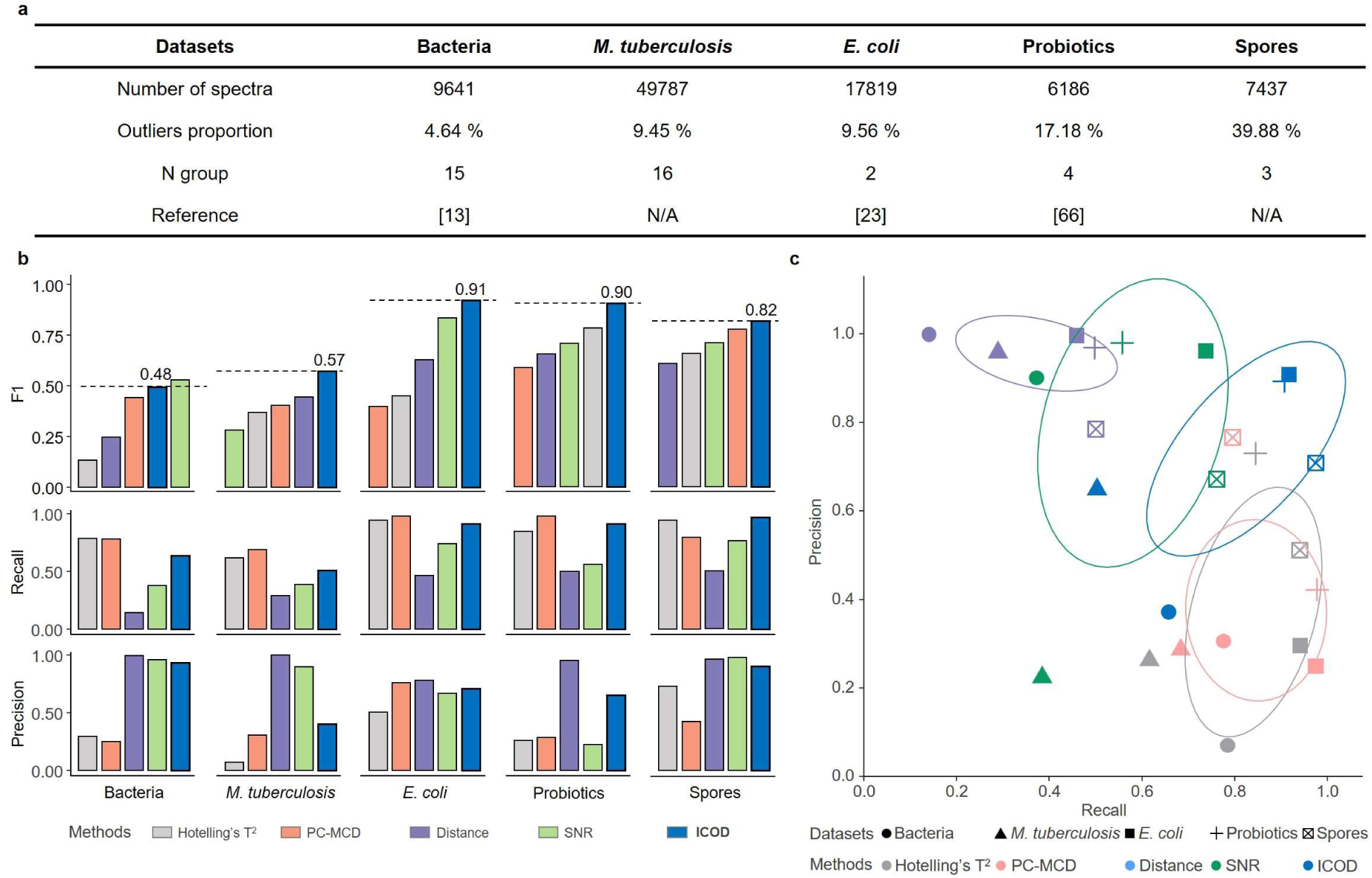
ICOD demonstrates superior performance over other methods on five public datasets. **a**, Summary of the five public datasets (for detailed descriptions, see **Methods**). **b**, F1 scores, recalls and precisions of five outlier detection methods on real datasets, the ground truth served as an artificial, item-by-item determination. **c**, Scatter plot showing the precision and recall value for the five datasets, where ICOD is always at a higher level than that obtained by other methods, demonstrating its superior performance and robustness.

### RamEx supports high-resolution single-cell metabolic profiling of microbial communities

To demonstrate the down-stream functionalities of RamEx, we present an analysis of spectral datasets collected at different stages of *Saccharomyces cerevisiae* pure culture fermentation, showcasing its advantages in analyzing single-cell metabolic within a population, with a particular attention to the impact of quality control on these functionalities. RamEx seamlessly integrates multiple non-linear dimensionality reduction algorithms and unsupervised clustering techniques, incorporating pre-dimensionality reduction and feature standardization specifically designed to address the collinearity and subtle variations inherent in scRaman datasets. To ensure efficient handling of large-scale integrated datasets, these methods are optimized through parallel computing and sparse processing. UMAP results revealed distinct structures associated with culture time (**Fig. 4a**), underscoring the effectiveness of RamEx in capturing dynamic biological changes. The Louvain method further grouped similar cell profiles into four distinct clusters based on ramanome-derived phenotypes (**Fig. 4b**). The formation and progression of these clusters correlate with the fermentation process, illustrating RamEx’s capability to effectively capture single-cell heterogeneity during fermentation as represented by the single-cell Raman spectrum. Subsequent composition analysis supported by RamEx elucidated the spatiotemporal composition of each defined cluster and highlighted the dynamic changes in cluster composition throughout the fermentation process (**Fig. 4c**).

**Figure 4.**
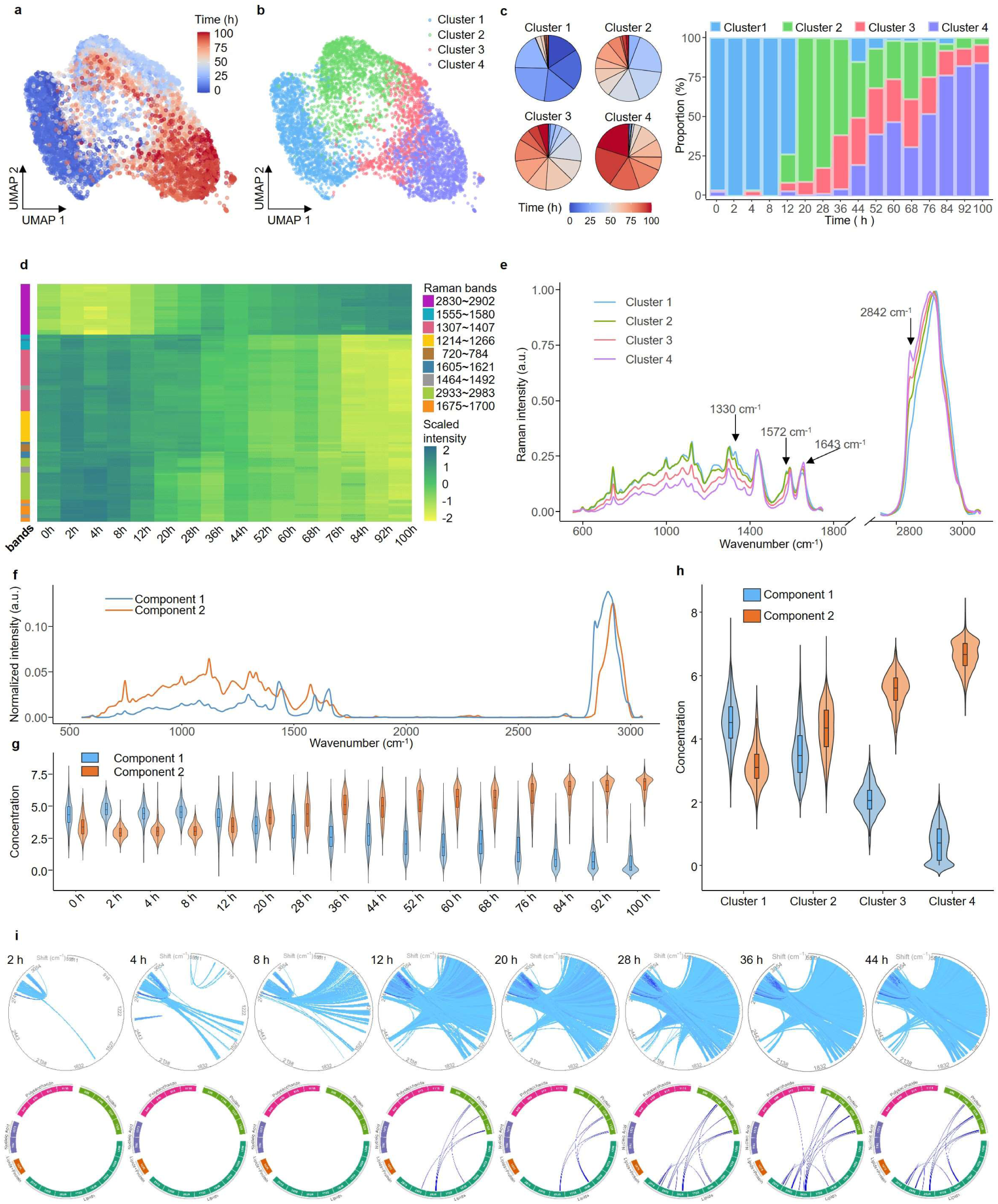
Application of RamEx to ramanomes from yeasts reveals intracellular substance changes over fermentation time. **a** and **b**, Uniform Manifold Approximation and Projection (UMAP) reduction of all ramanomes, colored by fermentation time or unsupervised clustering results. **c**, Time point composition of each cluster and cluster composition at each time point. Cluster 1 only appears in the first 12 h, while cluster 4 emerges after 36 h and increases over time. **d**, Heatmap of single markers shows the Raman intensity trends over time, identified from single-cell spectra. **e**, Mean spectra of 4 clusters, recorded by Raman markers derived from population-averaged spectra. **f** and **g**, Pure component spectral profiles and the corresponding concentration profiles resolved by MCR-ALS, though their concentrations show a more significant trend with cluster variation in (**h**). **i**, Global-IRCN and local-IRCN at each time point, with connections between Raman bands colored by their correlations.

Raman spectroscopy data encompass thousands of dimensions, many of which may not offer significant biological insights. Effective feature selection, aimed at identifying meaningful Raman markers, is essential for interpreting target phenotypes and providing focused information for subsequent experiments/sorting. RamEx perform marker identification by exploiting coherence among physically adjacent spectral bands and presents two distinct strategies: one focusing on single-cell spectra and the other on population-averaged spectra. Notably, RamEx identifies not only individual spectral bands but also paired combinations (e.g., *I_i_*/*I_j_*) that can function as Raman markers, enriching insights into the prior labeling and the functionality of each cell.

The single-cell spectral correlation-based approach is designed to uncover Raman bands that are associated with continues sequential variable. Specifically, RamEx identified Raman markers that associated with fermentation time, effectively divided them into three distinct temporal change types: gradual increases, gradient decreases, and sharp drops (**Fig. 4d**). This method, while effective in identifying patterns, contends with significant heterogeneity among individual cells, resulting in substantial overlap between different time points (**Supplementary Fig. 4b, c)**. The paired approach, however, mitigates this overlap to some extent (**Supplementary Fig. 4d, e**). Conversely, the population-averaged spectra method addresses baseline shifts caused by variations in adjacent peaks, efficiently eliminating spectral regions lacking chemical relevance. This method adeptly identifies emerging (e.g., 1643 cm⁻¹, 2842 cm⁻¹) and diminishing (e.g., 1330 cm⁻¹, 1575 cm⁻¹) peaks throughout the fermentation process (**Fig. 4e**, **Supplementary Fig. 5**). Such analysis provided by RamEx elucidates biological trends like the accumulation of fatty acids and triacylglycerides (TAGs) and the decrease in protein and nucleic acid levels, with clearer variations between clusters, thus shedding light on the biological rationale of cell clustering.

Raman spectroscopy inherently involves complex spectral bands, where different spectral bands may correspond to the same molecule, and many molecules exhibit similar trends. This complexity necessitates a spectral-level summary to accurately capture molecular changes. The “spectra decomposition” module in RamEx, which is distinct from the “Raman markers” module in RamEx, provides methods to unmix the spectra of intracellular mixtures. This unsupervised approach analyzes changes in molecular composition within single cells, focusing on alterations in macromolecular assemblies within a given ramanome. RamEx employs sophisticated methodologies by treating each spectrum as a linear combination of component spectra, enabling the decomposition of the spectral matrix into a component spectra matrix and a component concentration matrix. This approach allows RamEx to efficiently disentangle complex spectral data into interpretable components. Specifically, in the yeast dataset, MCR-ALS decomposes cell spectra into well-defined components linked to specific biomolecules— one highlighting lipid concentrations and another associated with other cellular substances (**Fig. 4f**). The strength of RamEx lies in its ability to reveal nuanced trends in concentration changes across fermentation timeframes (**Fig. 4g**). Despite the observed overlap, these decomposition techniques accentuate distinct concentration variations between different clusters (**Fig. 4h**), offering significant insights into the evolution of subpopulations within the fermentation system. Throughout diverse biological processes, phenotypic variations among individual cells exhibit continuous and dynamic patterns. RamEx offers Intra-Ramanome Correlation Network, which is an algorithm that we introduced (IRCN [62]). IRCA is adept at uncovering phenotypic correlations from a single snapshot of any cellular population, leveraging the inherent metabolic heterogeneity to provide insights into the intricate relationships within the population. For instance, the global intra-ramanome correlation generated by RamEx becomes increasingly intricate, indicating more active intracellular metabolite transformations (**Fig 4i, top**). Additionally, the local-IRCN focuses on changes at specific peaks rather than across all spectral bands, providing a more nuanced understanding of specific metabolite conversion (**Fig 4i, bottom**). By capturing these detailed intra-population dynamics from a single population, RamEx overcomes the limitations associated with traditional experimental gradients, emphasizing the importance of individual developmental trajectories in diverse biological systems.

Lately, we emphasized the critical importance of quality control by re-evaluating all downstream analyses using alternative QC methods in place of ICOD. The results demonstrated that RamEx significantly enhanced the clarity of phenotypic patterns by reducing data variance and improving the separation between different time points (**Supplementary Fig. 6-7**). ICOD proved to be particularly effective in maintaining the integrity of biological conclusions, offering a more consistent and reliable dimensionality reduction compared to prior approaches (**Supplementary Fig. 8-9**). Furthermore, the anticipated increase in IRCN activity with fermentation time was consistently tracked, reflecting predictable potential metabolite conversions (**Supplementary Fig. 10**). Spectral decomposition was notably improved, with MCR-ALS achieving successful convergence, thereby enhancing effectiveness of the decomposition process. These outcomes underscore the robustness of the quality control methods.

### RamEx enables high-throughput and scalable microbial ramanome analyses

Given that the number of single-cell individuals in microbial communities is typically enormous, RamEx incorporates a series of computational optimizations to enhance its efficiency in processing large datasets, including parallel computing frameworks, GPU acceleration, and C++ code optimizations. A key innovation is the use of optimized data structures and sparse matrix representations, which expedite computations in modules such as dimensionality reduction and clustering without compromising accuracy. Additionally, RamEx employs parallelization techniques at both the data and task levels, effectively distributing computational workloads across multi-core processors.

To evaluate the impact of these optimizations, we tested RamEx on an extensive dataset comprising approximately 270,000 spectra collected from 344 strains across four microbial species, encompassing over a thousand experimental batches (**Fig. 5a, b**). This dataset presented significant computational challenges due to its size, heterogeneity, and intrinsic complexity, serving as a robust benchmark for assessing computational efficiency and accuracy.

**Figure 5.**
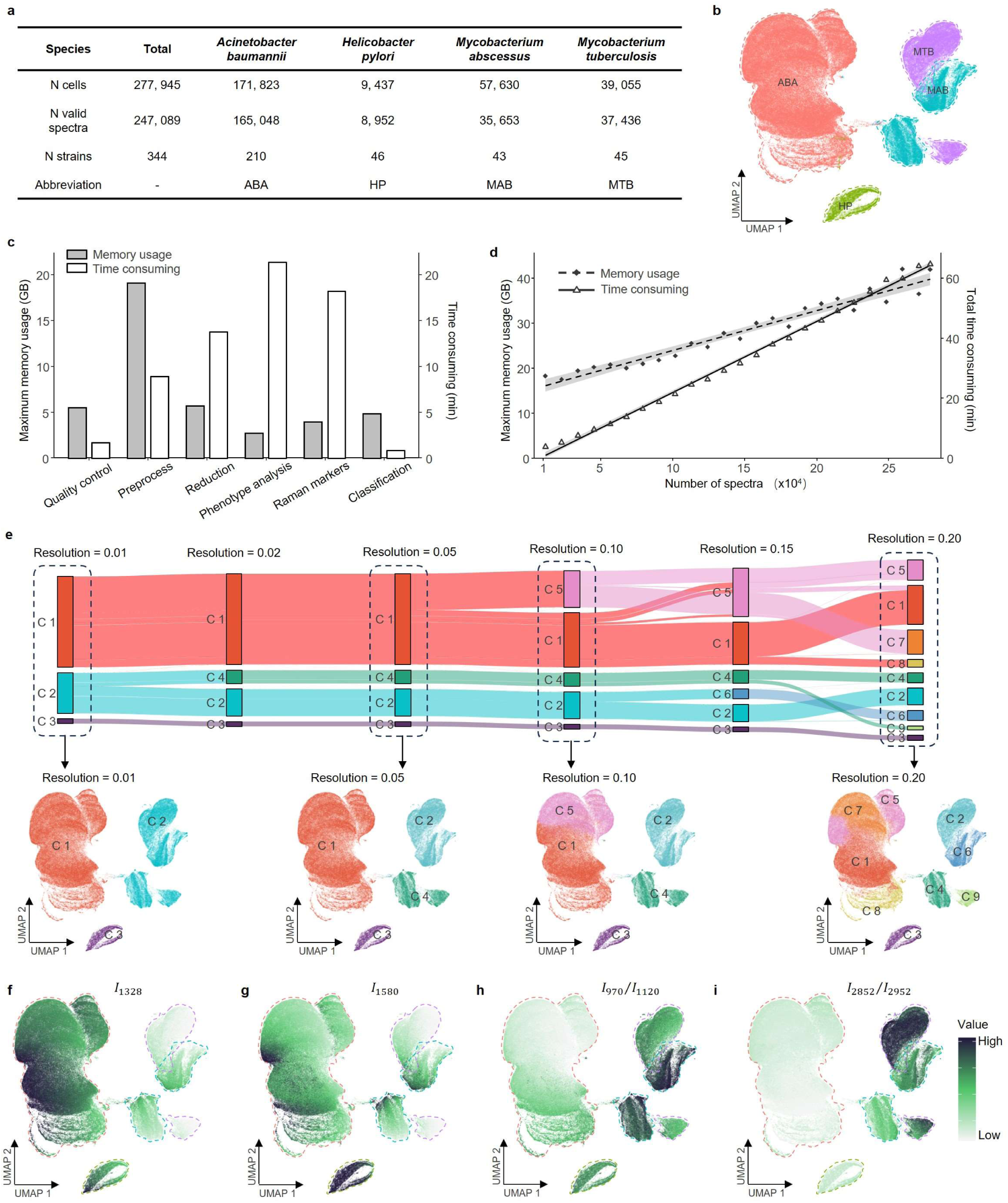
Application of RamEx to a large-scale ramanome dataset demonstrates high throughput and ability to unravel subtle yet important biological patterns. **a**, Basic information about the test dataset (details in **Methods**). **b**, UMAP projections colored by species. **c**, Runtime and Maximum memory usage of each module in RamEx. **d**, Total runtime and maximum memory usage with data volume. **e**, UMAP projections colored by the multi-level phenotype analysis with varying clustering resolution (top), with detailed community structures shown in the bottom projections. **f∼i**, UMAP projections colored by intensity or ratio values, representing specific Raman markers for ABA, HP, MAB and MTB, respectively.

RamEx substantially reduces computational time and resource consumption as compared to traditional methods. Specifically, the pipeline processed a 270,000-spectra dataset within 1 hour, while maintaining modest memory overhead (**Fig. 5c**). Scalability assessments across varying dataset sizes (from 10,000 to 270,000 spectra) revealed sub-linear scaling in both computational time and memory requirements (**Fig. 5d**), with individual modules exhibiting comparable performance characteristics (**Supplementary Fig. 11**). The same pipeline tackled the challenge on a much larger dataset of about 800,000 spectra, further illustrating its adaptability to large-scale data sets (**Supplementary Fig. 12**).

Firstly, we assessed the impact of quality control (QC) in ensuring data integrity. Outliers and low-quality spectra introduced excessive variability, obscuring genuine biological variations. For example, in PCA, outliers can skew principal components, masking true spectral features (**Supplementary Fig. 13**). Without sufficient QC, the “Raman markers” module’s ROC method failed to identify species-related tags effectively, as noises and outliers conceal critical spectral characteristics. QC methods significantly influence classification performance. Although SVMs are relatively robust to outliers, QC choice impact model accuracy and training efficiency. Among the evaluated methods, the ICOD method excelled, achieving a 99.87% prediction accuracy on the validation dataset, highlighting its effectiveness in retaining informative spectra while excluding aberrant data (**Supplementary Fig. 14a**). Conversely, overly stringent QC methods like Hotelling’s T^2^, PC-MCD, and SNR filtering retained only about 60% of the data, leading to resource wastage and potential loss of valuable information (**Supplementary Fig. 14b**). Distance-based methods also extended model training times, indicating slower SVM convergence due to retained noises (**Supplementary Fig. 14c**). Thus, by integrating ICOD, RamEx ensures both accurate and computationally efficient data analyses, which is crucial for large-scale datasets.

Secondly, the comprehensive QC process, combined with the capacity to analyze large-scale datasets, enables RamEx to uncover subtle biological patterns and relationships that are often undetectable in smaller datasets. For example, the enhanced “Phenotype analysis” module leveraged such high-quality, large-scale data to perform high-resolution clustering and visualization. The resulting phenotypic landscape captured the full spectrum of metabolic diversity across strains and species, illustrating subtle variations and transitions between phenotypic states (**Fig. 5e**). By varying the clustering resolution, we generated a multi-level data summarization that captures cell affiliations across adjacent resolutions (**Fig. 5e, top**). In this visualization, the thickness of the band segments is proportional to the number of cells, revealing phylogenetic relationships among species. Also, RamEx provides hierarchical clustering and drill-down functionalities, allowing users to navigate from broad overviews to detailed inspections of specific subpopulations within the dataset (**Fig. 5e, bottom**).

Finally, by analyzing the entire dataset collectively, the “Raman markers” module of RamEx uncovered unique Raman spectral features associated with specific species, contributing to a deeper understanding of their molecular compositions and metabolic processes. The ROC method (one-versus-rest receiver operating characteristic) was used to identify species-specific Raman bands and band combinations that achieve high area under the curve (AUC) values for classification tasks (**Supplementary Fig. 13**). Notably, *Acinetobacter baumannii* (ABA) and *Helicobacter pylori* (HP) exhibited higher signal intensities at 1328 cm⁻¹ and 1580 cm⁻¹, respectively, than other species (**Fig. 5f, g**). Additionally, peak ratio combinations such as *I*_2852_⁄*I*_2952_ and *I*_1000_⁄*I*_1078_ effectively differentiated *Mycobacterium abscessus* (MAB) from *Mycobacterium tuberculosis* (MTB) (**Fig. 5h, i**). These species-specific Raman markers provide higher-resolution molecular insights into each organism and demonstrate RamEx’s capability in handling complex classification tasks in large datasets. This capability offers algorithmic support for high-throughput identification and isolation of target cells.

## Discussion

RamEx was developed to address critical challenges in large-scale microbial ramanome analyses, offering an integrated solutions from spectral quality control, microbial phenotype classification, and advanced modeling across diverse biological contexts (**Fig. 1**). Key strength of RamEx lies in three components: robust outlier detection for designed microbial Raman spectra, specialized statistical and machine learning algorithms for spectral interpretation, and computational optimizations that enhance scalability and efficiency for high-throughput workflows.

A major challenge in high-throughput microbial Raman spectroscopy is the presence of anomalous spectra resulting from various acquisition artifacts, including empty acquisitions, misfocused spots, and device noises [50]. Such anomalies can obscure microbial spectral signatures and introduce biases in downstream phenotypic and classification analyses. Existing outlier detection methods often rely on empirical thresholds, limiting their adaptability in diverse microbial communities[55, 57, 59]. RamEx integrates ICOD to overcome this limitation by iteratively refining feature representations through peak-centric strategy instead of dimensionality reduction, thus preserving subtle yet essential spectral features. By adopting a peak-centric strategy and sharpening convolution, ICOD can accurately recognize clusters of anomalous spectra, achieving an F1 score of 0.97 even in datasets with up to 70% outliers (**Figures 2-3**, **Supplementary Fig. 2-3**), significantly surpassing the typical F1 scores of existing methods by at least 19.8% under real-world conditions. This parameter-free design preserves critical biological signals, circumventing inflated variance or spurious clusters in downstream analyses. In our validation tests, ICOD reduced the variance distribution along principal components from an outlier-induced extreme range (−1000 to 500) to a typical biological range, enabling clear cluster delineation and preserving meaningful spectral patterns (**Supplementary Fig. 6, 8–10**). Further benchmarking on extensive microbial datasets (totally over 90,000 spectra), spanning cell cultures and bacterial colonies, confirmed ICOD’s broad applicability across diverse microbial taxa with minimal need for hyperparameter adjustments (**Fig. 3, Supplementary Fig. 3**).

Beyond outlier detection, the microbial ramanome matrix exhibits distinct statistical and spectral characteristics that set it apart from other single-cell omics data. These include non-negative spectral intensities, non-linear molecular concentration-signal relationships, vibrational mode interactions, and wavelength band correlations. Additionally, systematic and random noise are inherent to Raman spectral data, further complicating computational analysis [67–69]. Such properties lead to statistical mismatches with conventional omics-optimized workflows, particularly those designed for sparse count matrices (e.g., scRNA-seq pipelines), making direct adaptation ineffective. To address these challenges, RamEx incorporates correlation-aware feature selection to handle spectral collinearity and implements non-negativity constraints in spectral decomposition, ensuring biologically meaningful spectral interpretations. Additionally, its robust preprocessing pipeline systematically removes noise while preserving key microbial spectral signatures. Unlike conventional omics tools that require extensive parameter tuning, RamEx provides empirically optimized default parameters, validated across diverse microbial datasets, ensuring optimal convergence and analytical rigor. By automating these complex processes, RamEx enhances the accessibility and reproducibility of large-scale microbial ramanome analyses, allowing biologists to extract reliable metabolic and phenotypic insights without requiring expertise in algorithmic fine-tuning.

Processing millions of spectra from diverse instruments requires both algorithmic efficiency and streamlined data management [70, 71]. To address these issues, RamEx integrates C++ computation with CPU+GPU parallel processing to achieve linear scalability (**Figure 5, Methods**) and standardizes data structures and output formats to enable reproducible workflows across acquisition platforms. Additionally, a Ramanome attribute table is provided for assessing spectral reproducibility, noise levels, and sample complexity (**Supplementary Fig. 1**), serving as a practical tool for initial data evaluation. While these efforts cannot fully resolve inter-instrument inconsistencies, RamEx establishes a standardized framework that enables researchers to share and analyze high-throughput Raman data across different laboratories and instruments, shedding light on systematic approaches to address multi-source spectral variability and improve reproducibility in microbial community analysis. Future developments may focus on implementing more robust batch correction methods, expanding libraries of pre-trained spectral models, and further automating hyperparameter tuning. As Raman flow cytometry becomes a routine tool for single-cell phenotyping, solutions like RamEx will be essential for delivering accurate, high-resolution insights into cellular metabolism.

## Conclusion

RamEx improves microbial ramanome analysis with its adaptive, threshold-free Iterative Convolutional Outlier Detection (ICOD), achieving exceptional spectral quality control (F1 > 0.97). By integrating modular workflows for phenotype clustering, metabolic fingerprinting, and intra-population heterogeneity analysis, it effectively bridges key computational gaps in Raman-based microbial phenomics. Optimized for scalability, RamEx efficiently processes million-scale datasets, enabling applications from antibiotic resistance tracking to functional dynamics studies at single-cell resolution. As a freely available R package, RamEx establishes a standardized framework for reproducible, high-throughput ramanomics, positioning Raman flow cytometry as a mainstream tool in microbial systems biology.

## Methods

### Data inputs

Five public datasets were used for evaluation: (*i*) “bacteria” dataset from Liu et al. [13], containing Raman spectra of 15 pathogenic bacterial isolates; (*ii*) “*Mycobacterium tuberculosis*” dataset from Mao et al., comprising single-cell Raman spectra of drug-sensitive and resistant MTB strains under various conditions; (*iii*) “probiotic” dataset from Zhang et al., containing spectra of four commercial probiotic strains collected using FlowRACS; (4) “*E. coli*” dataset from Wang et al. [23], including spectra of *E. coli* strains exposed to three antibiotics; and (*v*) “*Aspergillus candidus*” dataset of fungal spore spectra from wild-type, mutant, and edited strains collected via FlowRACS. Additionally, two self-collected datasets were used: a yeast dataset tracking *S. cerevisiae* during pilot-scale fermentation (16 time points, 0-100h), and a large-scale bacterial dataset comprising four species (*A. baumannii*, *H. pylori*, *M. abscessus*, and *M. tuberculosis*) across 344 strains under various experimental conditions. Both datasets were collected using a RACS-Seq instrument (Qingdao Single-cell Biotech, CN, 532 nm laser, 100 mW power, 1s integration, 100× objective).

### Data simulation

Simulated Ramans spectra were generated by Gaussian process regression (GPR) using the real Raman spectral data of *Escherichia coli* ATCC 25922 as reference [23]. Single-cell heterogeneity was simulated using radial basis function and white noise kernels, while instrumental variations were modeled by adding random Gaussian noise and polynomial baselines (orders 1-4, coefficients ±0.01). Four types of spectral anomalies [55] were simulated using homemade scripts: (*i*) Laser-induced burning effect: Signal overflow with sudden intensity spikes (60,000 counts); (*ii*) Optical defocusing: Selective enhancement/attenuation of spectral regions identified by the bubble method; (*iii*) Device artifact: Random addition of 20 overlapping peaks (width 200-1,000 cm⁻¹, intensity 100-1,000 counts); (*iv*) Empty acquisition: Substrate signals modeled with a 700 cm⁻¹ Gaussian peak and instrumental noise. The four types of anomalies were randomly introduced into the generated spectra, thus creating 19 anomalous datasets with anomaly proportions ranging from 5% to 95%. Five replicates of such datasets were generated for downstream analysis.

### Iterative Convolutional Outlier Detection

The ICOD (Iterative Convolutional Outlier Detection) is specifically designed to identify diverse outliers in the ramanome data, leveraging an iterative, unsupervised framework to enhance robustness and precision in high-dimensional spectral data. The method integrates peak detection via a modified bubble algorithm, fuzzy convolution, and clustering analysis to systematically isolate anomalous signals. Following preprocessing, all Raman spectra are initially treated as valid, except those with missing values. The original high-dimensional dataset *X_n_*_×*m*_ is then transformed into a reduced representation 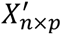 (where *p* ≪ *m*) through peak detection using a modified bubble method. This approach isolates critical spectral features while effectively discarding irrelevant noise and redundancy, ensuring the preservation of essential information for downstream analysis. Local anomalies are identified in the reduced dataset using the quartile method. For each peak *j*, an observation *x_ij_* is classified as a local outlier if it satisfies the conditions *x_ij_* < *Q*1*_j_* − 1.5 × *IQR_j_* or *x_ij_* > *Q*3*_j_* + 1.5 × *IQR_j_*, where *Q*1*_j_* and *Q*3*_j_* represent the first and third quartiles, respectively, and *IQR_j_* is the interquartile range. This process transforms the dataset into a Boolean matrix *B_n_*_×*p*_, where *B_ij_* = 1 if *x_ij_* is an outlier and *B_ij_* = 0 otherwise. The Boolean matrix is then subjected to k-means clustering with *k* = 2, separating spectra into normal and anomalous categories. To enhance the robustness of clustering, mean convolution filtering is applied to the Boolean matrix, reducing the influence of isolated local anomalies and improving the consistency of binary classification.

During this iterative process, a global outlier vector is iteratively refined, adjusting the probability of each spectrum being classified as an outlier. Higher values in the global outlier vector correspond to an increased likelihood of the spectrum being anomalous. The iterative refinement process recalibrates the classification by re-evaluating the spectra identified as normal spectrum in each cycle. This iterative process terminates when the global anomaly status stabilizes (i.e., no further updates occur) or when the proportion of anomalous bands falls below the predefined threshold.

### Performance evaluation of ICOD

The performance of ICOD and other outlier detection methods was evaluated using both simulated and real datasets. For the simulated datasets, the ground truth labels were generated alongside the random definition of anomalous spectra. For the real datasets, quality labels were manually assigned to each spectrum based on expert evaluation. To minimize bias from preprocessing, all data used for evaluation were processed using the default preprocessing pipeline of RamEx prior to outlier detection.

The evaluated methods included ICOD, the signal-to-noise ratio (SNR) method, the principal components - minimum covariance determinant (PC-MCD), the Hotelling’s T^2^ method, and the Euclidean distance method. The accuracy of quality control was assessed using three metrics: precision, recall, and F1 score. These metrics are defined as follows: 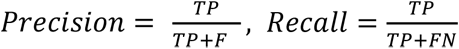, where True Positives (TP) represent spectra correctly identified as anomalous, False Positives (FP) represent spectra incorrectly classified as anomalous, and False Negatives (FN) represent anomalous spectra that were incorrectly classified as normal (undetected anomalies). F1 score is the harmonic mean of the precision and recall and is calculated as 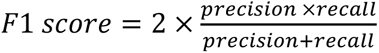. For the real datasets, the predicted labels generated by each of the five outlier detection methods were compared against the manually assigned ground truth labels, and the above evaluation metrics were calculated. For the simulated datasets, the metrics were computed for five parallel datasets with varying proportions of anomalies, providing a comprehensive assessment of each method’s performance under different conditions.

### Raman spectral preprocessing

The raw Raman spectra was preprocessed using the default pipeline in RamEx, including cosmic ray removal, spectral smoothing, baseline correction, and normalization. These steps were applied sequentially to ensure the quality and consistency of the spectral data. Cosmic ray removal was performed using a two-dimensional convolution-based spike detection method with a sharpening convolution kernel, where the center pixel value was set to 33. Signals in the identified cosmic ray regions were replaced with the corresponding intensities from adjacent spectra to minimizing distortion. Smoothing was conducted using the Savitzky-Golay method with a window size of 11 and a 5th-order polynomial fit, effectively reducing high-frequency noise while preserving spectral features. Baseline correction was then applied to remove low-frequency spectral noise using partitioned polynomial fitting. Specifically, a first-order polynomial was fitted to the 500–1800 cm⁻¹ region, while a sixth-order polynomial was applied to the 1800–3050 cm⁻¹ region to account for the varying baseline characteristics across the spectrum. Each spectrum was normalized to the maximum intensity value within the CH stretching region (2750–3050 cm⁻¹), ensuring comparability across spectra.

### Classification

A support vector machine (SVM) with linear kernel was employed for classification, using either a predefined test set or a 7:3 train-test split. To address the computational complexity of *O*(*d_L_N_S_*), where *N_S_* is the number of support vectors and *d_L_* is the dimensionality, PCA was applied to reduce the spectral data to 20 principal components before SVM training. Multi-class classification was handled using the “one-against-one” strategy, with test data projected into the same PCA space. The workflow’s performance was evaluated across different training set sizes. Using 10,000 randomly selected samples as a fixed test set, training sets ranging from 10,000 to 260,000 spectra were sampled from the remaining data. Both sets underwent ICOD and spectral preprocessing independently, with classification accuracy assessed on valid test spectra for each training size configuration.

### Marker Identification

Raman markers were identified as individual bands or band pairs strongly associated with ground truth labels. For continuous labels, pearson correlation coefficients were used, while for categorical labels, the area under the ROC curve (AUC) in a “one-versus-rest” framework quantified discriminative ability. Markers were selected when their correlations or AUC scores exceeded predefined thresholds. To overcome the computational complexity of evaluating band combinations 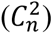, the analysis was accelerated using C++ implementation via “Rcpp” with efficient memory management and multi-core parallelization. While RamEx defaults to single-band analysis, this study evaluated both single-band and paired-band combinations for comprehensive marker identification.

### Feature reduction and phenotype analysis

Nonlinear dimensionality reduction methods, such as Uniform Manifold Approximation and Projection (UMAP) and t-distributed Stochastic Neighbor Embedding (t-SNE), were employed for the visualization of ramanome datasets. To ensure computational efficiency, the spectral matrix was first subjected to principal component analysis (PCA) for preliminary dimensionality reduction. Additionally, the visualization process was accelerated using multithreaded computation. Cellular phenotypes within the ramanome were identified using the Louvain community detection method based on shared nearest neighbor (SNN) modularity optimization. After computing k-nearest neighbors and constructing an SNN graph using cosine similarity, clusters were identified through modularity optimization. Small clusters (<1%) were merged with their nearest communities, and for datasets exceeding 10,000 samples, the SNN graph was constructed based on sparse distance matrix. Multiple resolution parameters (0.01-0.2) were applied to generate a hierarchical phenotypic structure, with results visualized using UMAP.

### Spectral decomposition

To decompose the dataset into its underlying components and corresponding concentration profiles, the Multivariate Curve Resolution-Alternating Least Squares (MCR-ALS) algorithm was employed. The algorithm iteratively optimizes the decomposition of the dataset *X* into two matrices, as described by the following equation:

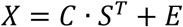

where *C* represents the concentration matrix, *S* is the spectral matrix, and *E* denotes the residual matrix. Nonnegativity constraints were imposed on both concentration and spectral components. Additionally, an angle constraint was applied to minimize collinearity between components during the optimization process.

### Intra-ramanome analysis

Intra-ramanome correlation analysis (IRCA) was conducted to observe the heterogeneity within the ramanome and to elucidate intracellular metabolite conversions. For detailed methodological descriptions, refer to He et al. [62]. In this study, global and local intra-ramanome correlation networks (global-IRCNs and local-IRCNs) were constructed at different fermentation time points, with thresholds of *p* < 0.05 and *ρ* < −0.6. The global-IRCN considers variations across all Raman bands, providing a comprehensive overview of the ramanome’s dynamic changes, whereas the local-IRCN focuses on specific bands of interest that represent key biomolecular components. For example, bands at 781 and 1573 cm^−1^ correspond to nucleic acids; 847, 886, 915, and 1117 cm^−1^ to polysaccharides; 995, 1148, 1235, and 1328 cm^−1^ to proteins; and 1295, 1438, 1591, 1647, 1738, 2840, 2900, and 2919 cm^−1^ to lipids. Additionally, the band at 1075 cm^−1^ reflects a combination of lipid and protein signals.

### Memory usage and time-consuming tests

Tests on a large dataset containing Raman spectra from four bacterial species were used. Random subsets of 1,000 to 270,000 spectra were sampled and analyzed following the RamEx workflow, including six main steps: data import, quality control, spectral preprocessing, feature reduction, Raman marker identification, and classification. For classification, the dataset was split into training and testing sets with a 7:3 ratio. The time consumption of each step was recorded using R’s built-in “Sys.time()” function to measure the intermediate processing times. Peak memory usage during the analysis was monitored and measured using the “Rprof” function to capture the maximum memory allocation. All analyses were performed on a CentOS Linux 7 server with an Intel Xeon E7-4820 v4 processor (80 cores, 2.00 GHz), 575 GB RAM, using R 4.3.0 and OpenBLAS 0.3.3 with LAPACK 3.8.0.

## Abbreviations

ABA: Acinetobacter baumannii
AUC: Area Under the Curve
GMM: Gaussian Mixture Model
GPU: Graphics Processing Unit
HP: Helicobacter pylori
ICOD: Iterative Convolutional Outlier Detection
ICA: Independent Component Analysis
IRCA: Intra-Ramanome Correlation Analysis
IRCN: Intra-Ramanome Correlation Network
LDA: Linear Discriminant Analysis
MAB: Mycobacterium abscessus
MCR-ALS: Multivariate Curve Resolution–Alternating Least Squares
MLR: Multiple Linear Regression
MTB: Mycobacterium tuberculosis
NMF: Non-negative Matrix Factorization
PC-MCD: Principal Components–Minimum Covariance Determinant
PCA: Principal Component Analysis
PLS: Partial Least Squares Regression QC Quality Control
RBCS: Raman Barcode of Cellular-response to Stresses
ROC: Receiver Operating Characteristic
SCRS: Single-Cell Raman Spectroscopy
SNR: Signal-to-Noise Ratio
SVM: Supporting Vector Machine
t-SNE: t-distributed Stochastic Neighbor Embedding
TAGs: Triacylglycerides
UMAP: Uniform Manifold Approximation and Projection

## Data availability

All datasets that support the findings of this study have been deposited in ScienceDB and can be accessed from https://www.scidb.cn/en/s/ju6VNf. Moreover, additional larger-scale dataset was downloaded from Hill I. E. et al.[72]

## Acknowledgements

We are grateful for Fengyun Lv’s assistance in simulating data generation and thank Lei Zhang for the insightful discussions on software design.

## Funding

This work was supported by grants from National Natural Science Foundation of China (32030003), National Key Research and Development Program of China (2022YFF0713102), Key Technology Research and Development Program of Shandong Province (2024KJHZ033), Natural Science Foundation of Shandong Province (ZR2022QC020) and National Natural Science Foundation of China (32200448).

## Author information

Yanmei Zhang, Gongchao Jing contributed equally to this study.

## Contributions

L.S. and Y.Z. conceived and designed the studies; Y.Z., G.J, R.C. and Y.H. wrote codes in the software; X.W., J.Z. offered their published datasets for module tests; Y.L., Y.M. performed additional experiments for further exhibit RamEx’s performance; H.Y. collected these datasets; Y.Z. and G.J. prepared the tables and figures; Y.Z. and L.S. wrote the manuscript; L.S. and J.X. revised the manuscript; Y.G. and X.Z. provided procedural advice; G.J., Y.W. and M.W modified this software to make it complete.

## Corresponding authors

Correspondence to Luyang Sun or Jian Xu.

## Competing interests

The authors declare no competing interests.

## Notes

https://www.scidb.cn/en/s/ju6VNf

### Competing Interest Statement

The authors have declared no competing interest.

