## Supplementary tables and figures for "RamEx: An R package for high-throughput microbial ramanome analyses with accurate quality assessment"

**Supplementary table 1**. Currently available softwares for Ramanome analysis

| Name | Outlier detection | Biosample optimization | Speed optimization for big data | Pipeline for Ramaome analysis | GUI | Year | Reference |
| --- | --- | --- | --- | --- | --- | --- | --- |
| LabSpec | / | / | / | / | Y |  |  |
| WiRE | / | / | / | / | Y |  |  |
| WITec | / | / | / | / | Y |  |  |
| ChemoSpec | / | / | Limited | / | / | 2008 | [1] |
| HyperSpec | PCA | / | Limited | Y | / | 2011 | [2] |
| MALDIquant | / | / | / | / | / | 2012 | [3] |
| MCR-ALS | / | Y | Limited | / | Y | 2015 | [4] |
| Rametrix™ | / | Y | Partial | Y | Y | 2018 | [5] |
| NWUSA | / | Y | / | / | Y | 2020 | [6] |
| Rampy | / | / | Extendable | / | / | 2021 | [7] |
| CELL IMAGE | / | Y | / | Y | Y | 2022 | [8] |
| RamanSPy | / | Y | Y | Y | / | 2024 | [9] |

**Supplementary Table 2**. List of datasets used to test the availability of RamEx

| Datasets | Source | Species | Strain | Number of spectra | Device | Ref. |
| --- | --- | --- | --- | --- | --- | --- |
| Bacteria | Min Liu et.al | *Lactobacillus fermentum* | ATCC 9338 | 698 | CAST-R  #1 | [10] |
|  |  | *Staphylococcus epidermidis* | ATCC 12228 | 550 |  |  |
|  |  | *Rothia dentocariosa* | ATCC 17931 | 635 |  |  |
|  |  | *Enterococcus faecium* | ATCC 19434 | 592 |  |  |
|  |  | *Acinetobacter baumannii* | ATCC 19606 | 602 |  |  |
|  |  | *Escherichia coli* | ATCC 25922 | 588 |  |  |
|  |  | *Helicobacter pylori* | ATCC 26695 | 587 |  |  |
|  |  | *Pseudomonas aeruginosa* | ATCC 27853 | 641 |  |  |
|  |  | *Staphylococcus sciuri* | ATCC 29061 | 559 |  |  |
|  |  | *Enterococcus faecalis* | ATCC 29212 | 604 |  |  |
|  |  | *Staphylococcus aureus* | ATCC 29213 | 617 |  |  |
|  |  | *Staphylococcus capitis* | ATCC 49326 | 605 |  |  |
|  |  | *Enterobacter hormaechei* | ATCC 700323 | 573 |  |  |
|  |  | *Enterococcus casseliflavus* | ATCC 700327 | 585 |  |  |
|  |  | *Klebsiella pneumoniae* | ATCC 700603 | 582 |  |  |
| *M. tuberculosis* | Yuli Mao et.al | *Mycobacterium tuberculosis* | ATCC 27294 (drug-sensitive) | 11562 | CAST-R  #2 |  |
|  |  |  | 3 clinical isolates (drug-resistant) | 38225 |  |  |
| *E. coli* | Xixian Wang et.al | *Escherichia coli* | ATCC 25922 | 8903 | FlowRACS | [11] |
|  |  |  | clinical isolates 1768859 | 8916 |  |  |
| Probiotics | Jia Zhang et.al | *Lactiplantibacillus plantarum* | CICC 6009 | 1409 | FlowRACS | [12] |
|  |  | *Streptococcus thermophilus* | CICC 6063 | 1701 |  |  |
|  |  | *Lactobacillus reuteri* | BNCC 192190 | 1678 |  |  |
|  |  | *Bifidobacterium animalis* | BNCC 186305 | 1398 |  |  |
| Spores | Yanmei Zhang et.al | *Aspergillus candidus* | wild-type strain | 2518 | FlowRACS |  |
|  |  |  | high-yield chloroflavonin mutants | 2193 |  |  |
|  |  |  | genetically edited strain lacking chlorflavonin production | 2726 |  |  |
| Yeasts | Yuandong Li et.al | *Saccharomyces cerevisiae* | CICC 30018 | 9208 | CAST-R  #2 |  |
| Bacteria-4 species | Pengfei Zhu et.al | *Acinetobacter baumannii* | 108 clinical isolates | 171823 | CAST-R  #1  CAST-R  #2 | [10, 13, 14] |
|  | Min Liu et.al | *Helicobacter pylori* | 19 clinical isolates | 9437 |  |  |
|  | Yuli Mao et.al | *Mycobacterium abscessus* | 5 clinical isolates | 37599 |  |  |
|  |  |  | ATCC19977 | 20031 |  |  |
|  |  | *Mycobacterium tuberculosis* | 44 clinical isolates | 36846 |  |  |
|  |  |  | ATCC 27294 | 2209 |  |  |
| Human cells | Iona E. Hill et.al | UVW human glioma cells | synchronised | 430590 | Renishaw InVia | [15] |
|  |  |  | unsynchronised | 362717 |  |  |
| **Total** | | **24 species** | **201 strains** | **1,170,707 spectra** | **4 devices** |  |

**Supplementary table 3**. Functions contained in RamEx

| Module | Algorithm | Code | Description | Reference |
| --- | --- | --- | --- | --- |
| Basic Module |  | read.spec | Load data from diverse instrument platforms |  |
|  |  | Ramanome | Define a standardized format for Ramanomes |  |
| Quality control | Iterative Convolutional Outlier Detection (ICOD) | Qualitycontrol.ICOD | Detect outliers among all samples via sudden intensity changes in spectra |  |
|  | Minimum covariance determinant (MCD) | Qualitycontrol.Mcd | A highly robust estimator of multivariate location and scatter | [16] |
|  | Hoteling’s T^2^ | Qualitycontrol.T2 | A multivariate statistical test used to determine whether the mean of the two samples is significantly different | [17] |
|  | Euclidean distance | Qualitycontrol.Dis | The Euclidean distance is a measure of the straight-line distance between two points in Euclidean space | [18] |
|  | Signal-to-noise ratio (SNR) | Qualitycontrol.Snr | A measure used in science and engineering to quantify how much a signal has been corrupted by noise | [19] |
| Preprocesssing |  | Preprocesssing.Background.Remove | Substract solute background from Raman spectra |  |
|  |  | Preprocesssing.Background.Spike | Remove cosmic spikes from Raman spectra |  |
|  | Savitzky-Golay filtering | Preprocesssing.Smooth.Sg | General least-squares smoothing and differentiation by the convolution (Savitzky-Golay) method | [20] |
|  |  | Preprocesssing.Normalize | Normalize the by maximum intensity, intensity at specified wavelength or total area under the curve |  |
|  |  | Preprocesssing.Cutoff | Spectral truncation |  |
|  | Polynomial fitting | Preprocesssing.Baseline.Polyfit | Corrects baseline using polynomial fitting | [21] |
|  | Bubble | Preprocesssing.Baseline.Bubble | Corrects baseline using bubble method, more suitable for liquid phase acquisition spectra | [22] |
| Feature reduction | Uniform Manifold Approximation and Projection (UMAP) | Feature.Reduction.Umap | A nonlinear dimensionality reduction method that preserves global structure of the data by modeling it as a manifold | [23] |
|  | t-distributed Stochastic Neighbor Embedding (t-SNE) | Feature.Reduction.Tsne | A nonlinear dimensionality reduction technique that focuses on preserving local relationships by minimizing the divergence between high-dimensional and low-dimensional pairwise similarities | [24] |
|  | Principal component analysis (PCA) | Feature.Reduction.Pca | A linear dimensionality reduction method that transforms data into a set of orthogonal components, maximizing variance in the lower-dimensional space | [25] |
|  | Principal Coordinates Analysis (PCoA) | Feature.Reduction.Pcoa | Similar to PCA but works directly with distance matrices, making it suitable for non-Euclidean data | [26] |
|  |  | Feature.Reduction.Intensity | Extract peak intensities at specific wavelengths |  |
| Classification | Ramdom forest | Classification.Rf | An ensemble learning method that builds multiple decision trees and combines their outputs (e.g., majority voting for classification) | [27] |
|  | Linear discriminant analysis (LDA) | Classification.Lda | A linear classification method that projects data into a lower-dimensional space to maximize class separability, works better for linearly separable data and small datasets | [28] |
|  | Supporting vector machine (SVM) | Classification.Svm | Finds the optimal hyperplane to separate classes by maximizing the margin between them, powerful for complex, high-dimensional data but computationally expensive | [29] |
|  | Gaussian Mixture Model (GMM) | Classification.Gmm | A probabilistic model that assumes data is generated from a mixture of Gaussian distributions and assigns probabilities to each class | [30] |
| Quantification | Partial Least Squares (PLS) | Quantification.Pls | A regression method that reduces predictors to a smaller set of latent variables while maximizing the covariance between predictors and response variables, ideal for situations where predictors are highly collinear or when the number of predictors exceeds the number of observations | [31] |
|  | Multiple linear regression  (MLR) | Quantification.Mlr | Simple and interpretable regression method, but assumes no multicollinearity among predictors and a linear relationship between predictors and response | [32] |
|  | Generalized linear model (GLM) | Quantification.Glm | Extends linear regression by allowing the dependent variable to follow distributions other than normal (e.g., binomial, Poisson) and uses a link function to relate predictors to the response | [33] |
| Raman markers | Raman Barcode of Cellular-response to Stresses (RBCS) | Raman.Markers.Rbcs | Identifies important spectral regions using machine learning methods, bands with high contributions to the classifier are considered Raman barcodes | [34] |
|  | Receiver operating characteristic (ROC) curve | Raman.Markers.Roc | Identifies Raman markers by measuring the discriminative power of specific bands in distinguishing between given labels |  |
|  | Correlations | Raman.Markers.Correlations | Identifies Raman markers by measuring the linear relationships of specific bands, given labels should be meaningful continuous values |  |
| Phenotype analysis | Louvain | Phenotype.Analysis.Louvaincluster | A community detection algorithm primarily used for clustering in networks or graphs by maximizing modularity (a measure of cluster quality) | [35] |
|  | k-Means | Phenotype.Analysis.Kmeans | A centroid-based clustering algorithm that partitions data into a predefined number of clusters by assigning sample to the nearest center | [36] |
|  | Hierarchical clustering analysis (HCA) | Phenotype.Analysis.Hca | Builds a hierarchy of clusters by either merging smaller clusters (agglomerative) or splitting larger ones (divisive) based on a distance metric | [37] |
| Spectral decomposition | Multivariate Curve Resolution-Alternating Least Squares (MCR-ALS) | Spectral.Decomposition.Mcrals | Decomposes mixed spectra into pure component spectra and their concentrations using constraints (e.g., non-negativity) | [38] |
|  | Independent Component Analysis (ICA) | Spectral.Decomposition.Ica | A statistical method that separates mixed signals into statistically independent components by maximizing non-Gaussianity, effectively separating overlapping signals | [39] |
|  | Non-negative matrix factorization (NMF) | Spectral.Decomposition.Nmf | Decomposes data into non-negative components, ensuring that both the basis and coefficients are non-negative, with less constrains compared to MCR-ALS | [40] |
| Intra-Ramanome analysis | Intra-Ramanome Correlation Analysis (IRCA) | Intraramanome.Analysis.Irca.Global | Analyzes correlations between Raman bands within a Ramanome to identify co-varying spectral features | [41] |
|  |  | Intraramanome.Analysis.Irca.Local | Analyzes correlation between interested Raman bands record by defined attribution | [41] |
|  | 2D Correlation Spectroscopy Analysis (2D-COS) | Intraramanome.Analysis.2Dcos | Captures both synchronous (simultaneous changes) and asynchronous (sequential changes) relationships, providing detailed insights into spectral changes | [42] |

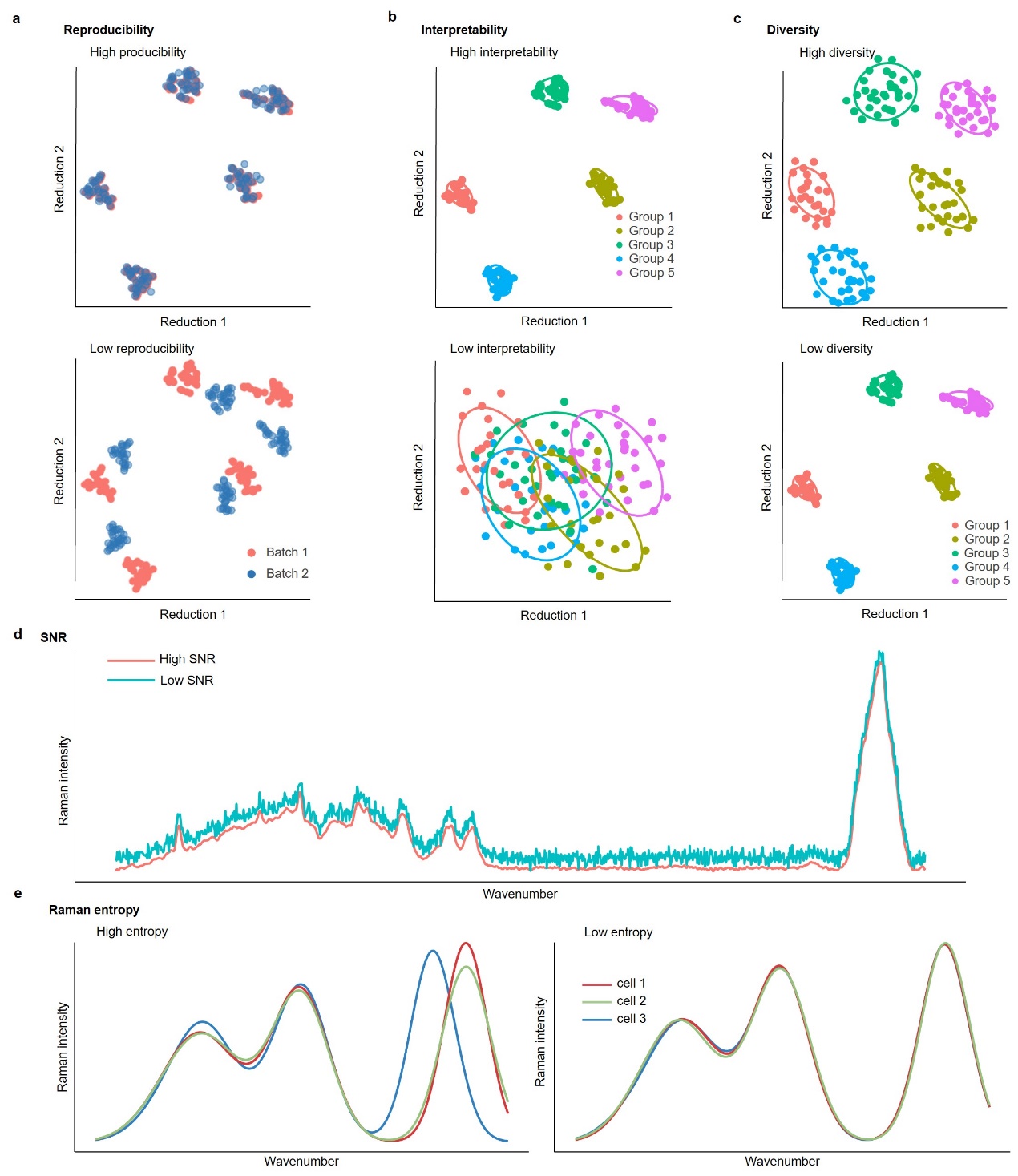

**Supplementary figure 1**. **Detailed descriptions for Raman Attribute Table**. **a**, Reproducibility measures the robustness of Raman features across batches by assessing spectral differences due to biological replicates and variations in Raman acquisition equipment. **b**, Interpretability refers to the proportion of variance in the spectral matrix explained by a specific metadata factor, indicating the impact of experimental factors on the spectra. **c**, Diversity denotes the proportion of variance within the Ramanome, reflecting substantial single-cell heterogeneity. **d**, Signal-to-Noise Ratio (SNR) is adapted from the traditional definition in compound spectra, specifically refined for biological spectra: the signal is defined as the highest intensity within the fingerprint region, and noise is the average intensity in silent regions. **e**, Raman Entropy reflects the Shannon entropy of peak positions and intensities, focusing on variations in Raman peaks rather than the entire spectrum, and describes total variance unrelated to metadata.

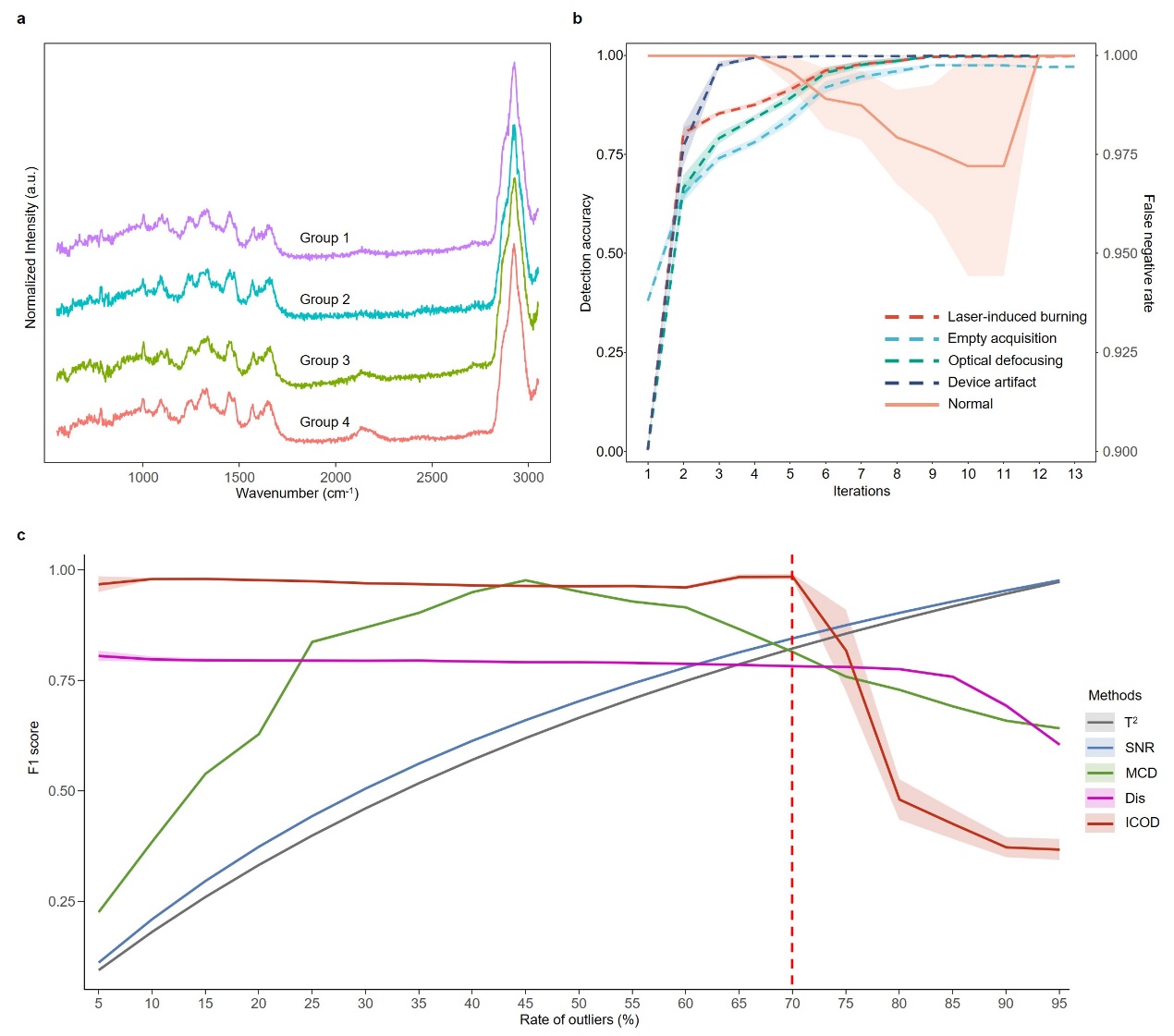

**Supplementary figure 2**. **Performance tests of outlier detection methods on simulated datasets**. **a**, Simulated spectra generated using a Gaussian Mixture Module. **b**, Detection rates of abnormal spectra or retention rates of normal spectra across ICOD’s iterations. **c**, F1 scores with the proportion of anomalies.

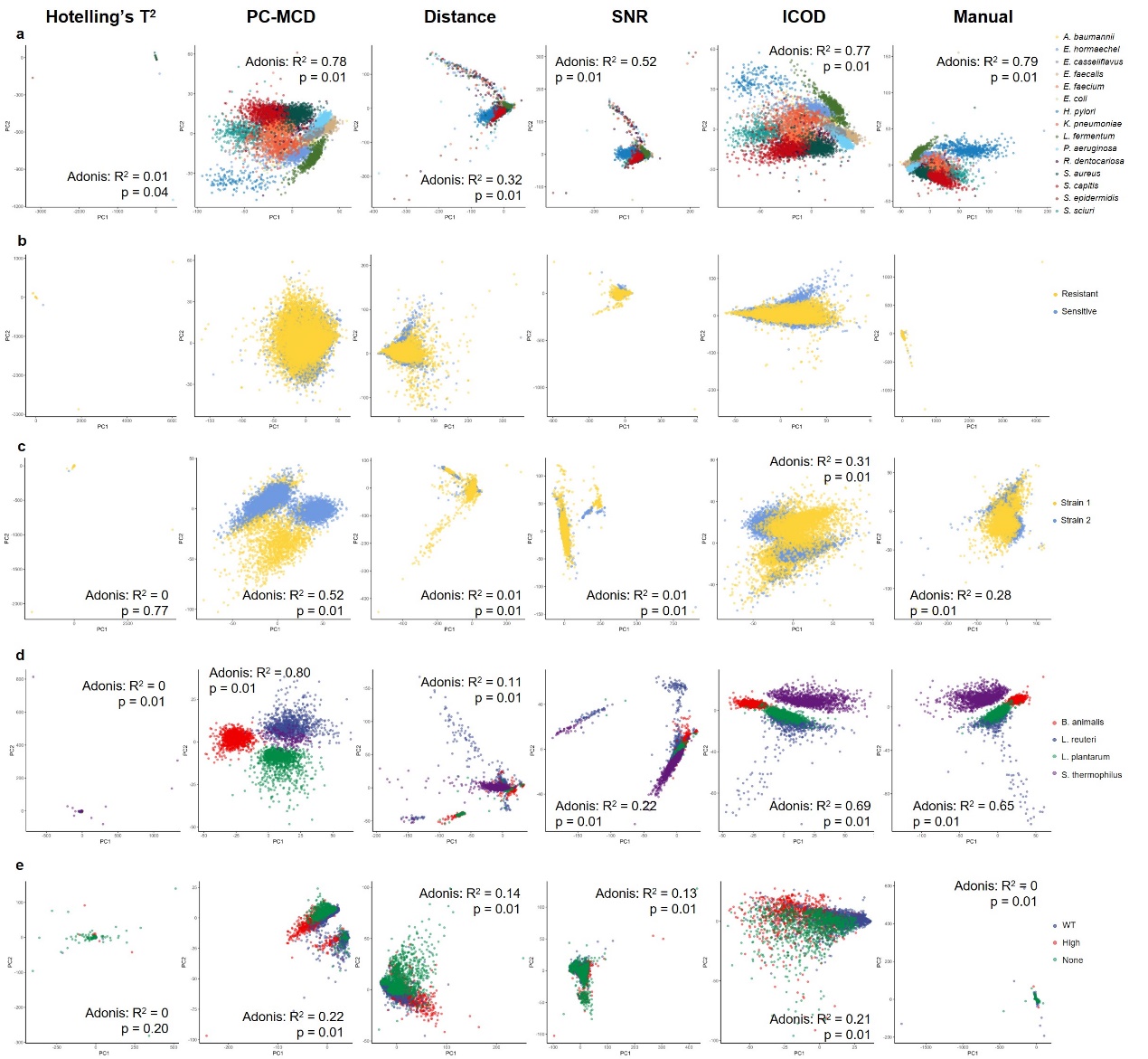

**Supplementary figure 3**. **PCA visualizations of five open-source datasets managed by five different methods**. **a**, “Bacteria” dataset. **b**, “*M. tuberculosis*” dataset. **c**, “*E. coli*” dataset. **d**, “Probiotics” dataset. **e**, “Spores” dataset.

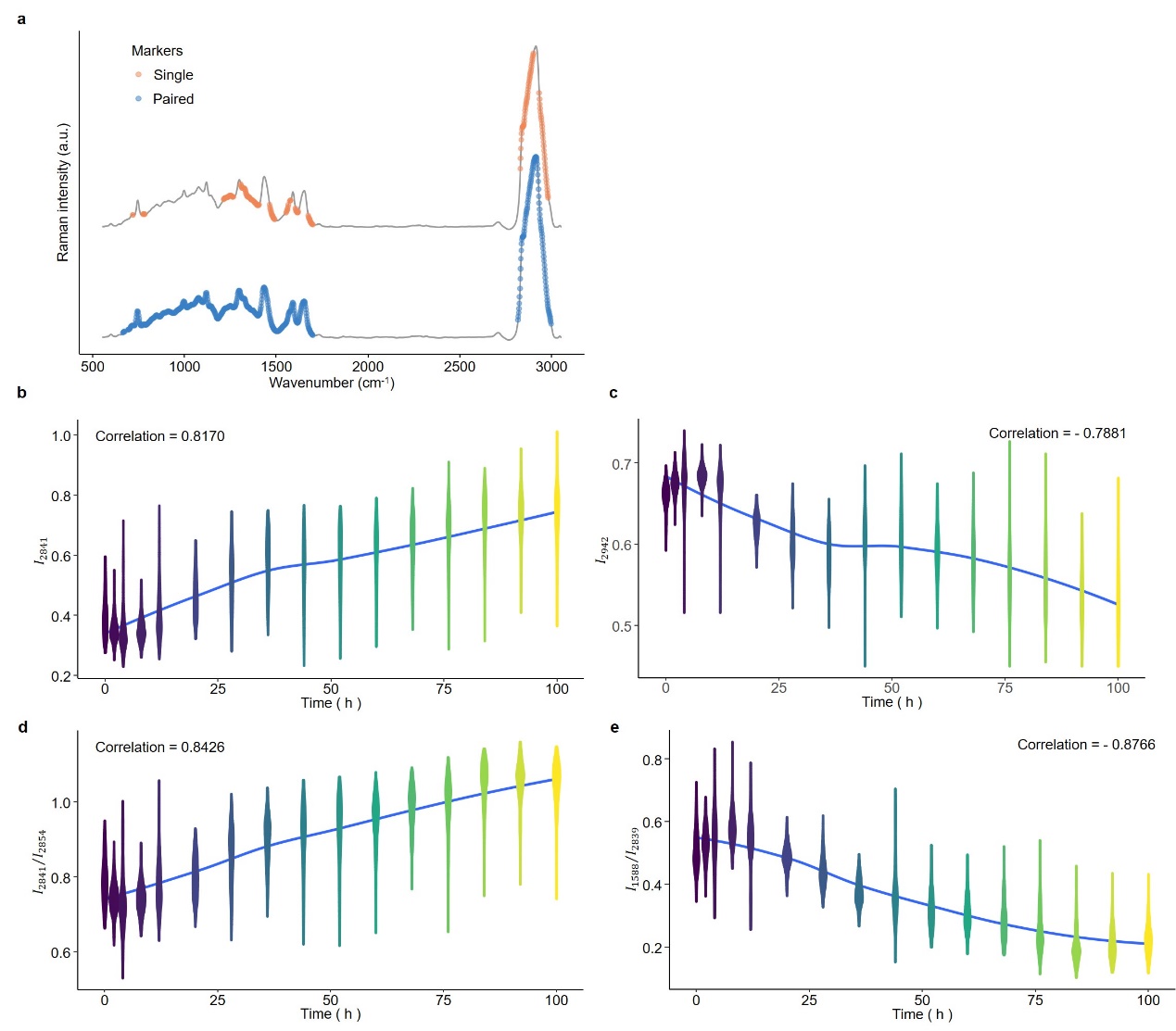

**Supplementary figure 4**. **“Raman markers” module in RamEx**. **a**, Selected markers identified using the “correlation” method. **b**, **c**, Singular Raman markers most associated with fermentation times, showing positive or negative correlations. **d**, **e**, Paired Raman markers most associated with fermentation times, showing positive or negative correlations.

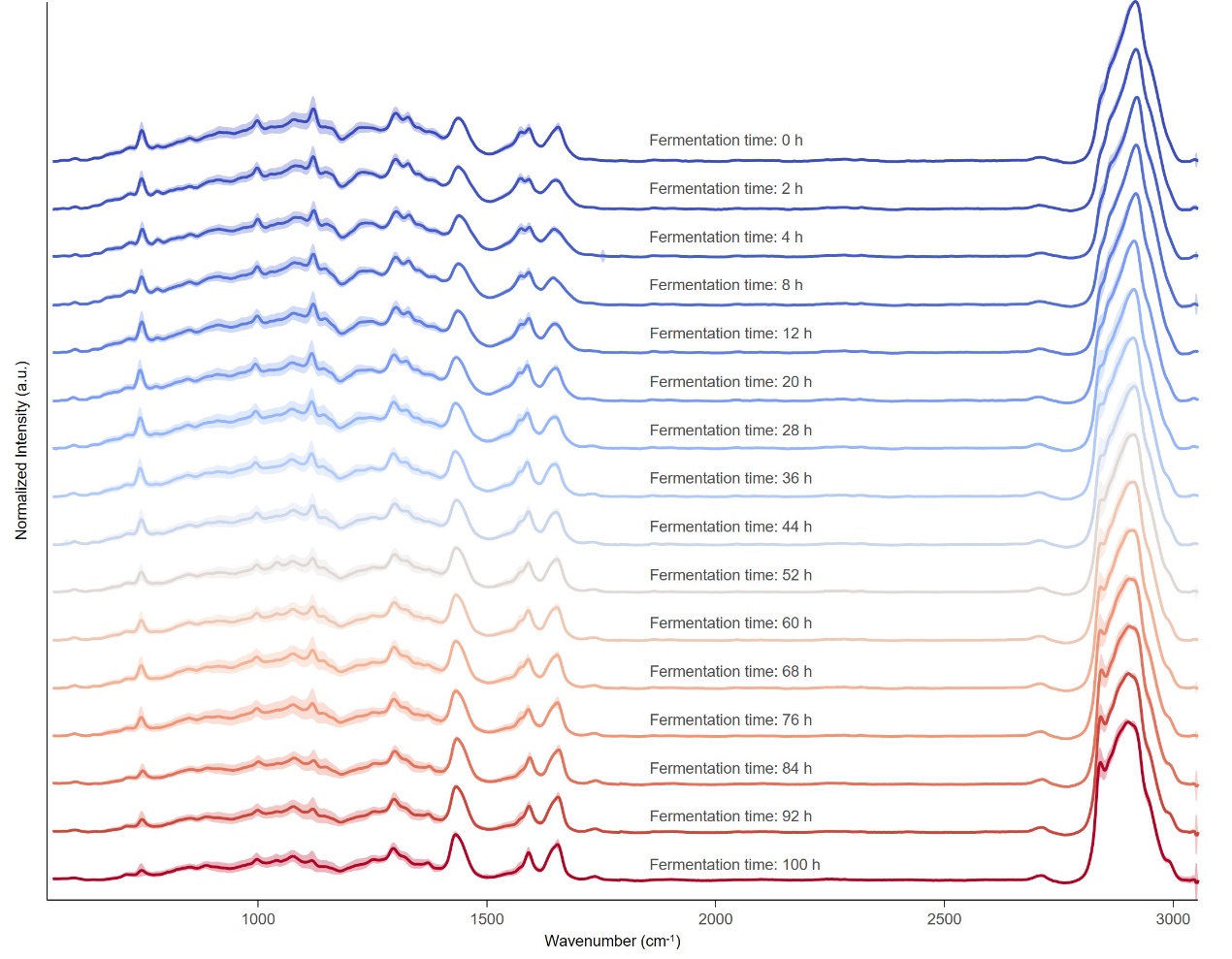

**Supplementary figure 5**. **Mean spectrum of each fermentation time point**. That is hard to find Raman markers by visual observation.

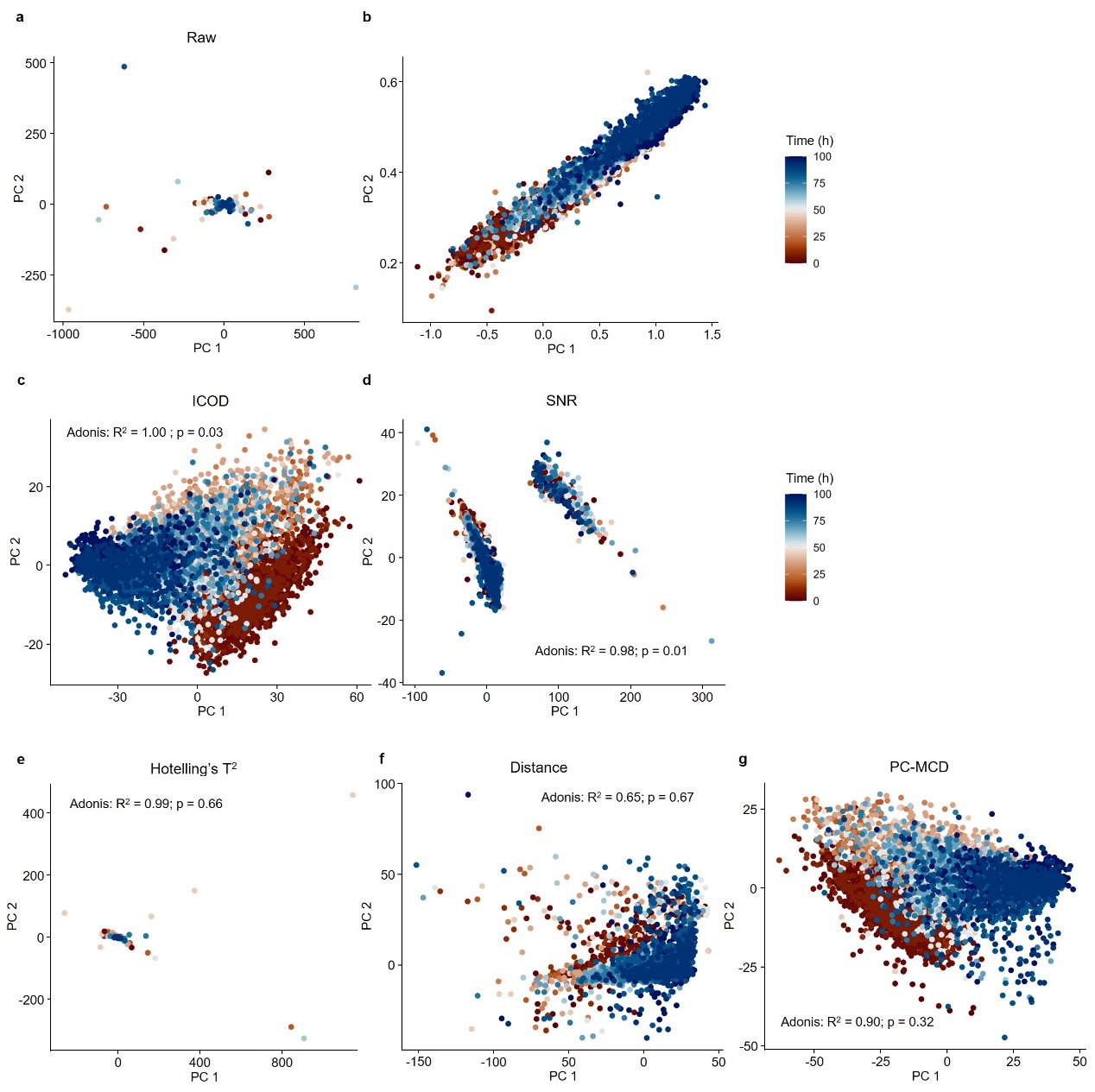

**Supplementary figure 6**. **PCA visualizations of the yeast dataset managed by five different methods**. **a**, Reductions of the uncontrolled dataset, exhibiting extreme variance ranges from -1000 to 500. **b**, Normal spectra are clustered in a narrow range of -1.0 to 1.5. **c**, Data matrix controlled by the ICOD method, showing a clear separation between fermentation times (Adonis: $R^{2}=1.00, p=0.03$). **d**, Data matrix controlled by SNR method, with samples divided into two groups. **e~g**, Data matrix controlled by Hotelling’s T^2^, Euclidean distance, and PC-MCD, respectively.

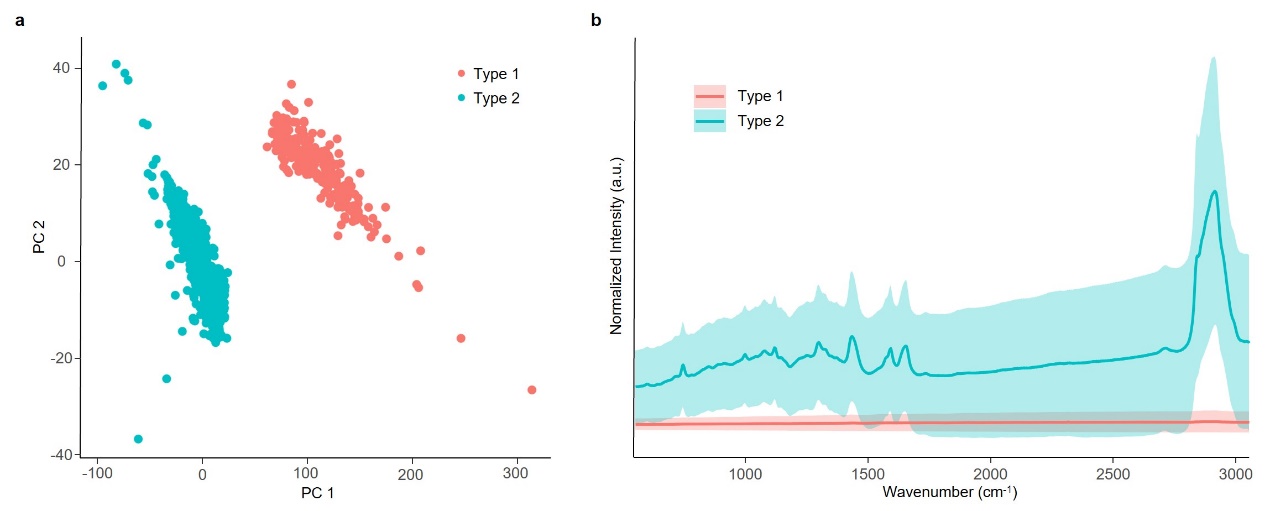

**Supplementary figure 7**. **Two spectral types in the data matrix managed by the SNR method**. **a**, Two distinct data types in the “valid” spectra. **b**, Mean spectra of these two types, with “type 2” mainly consisting of the spectra from null acquisition.

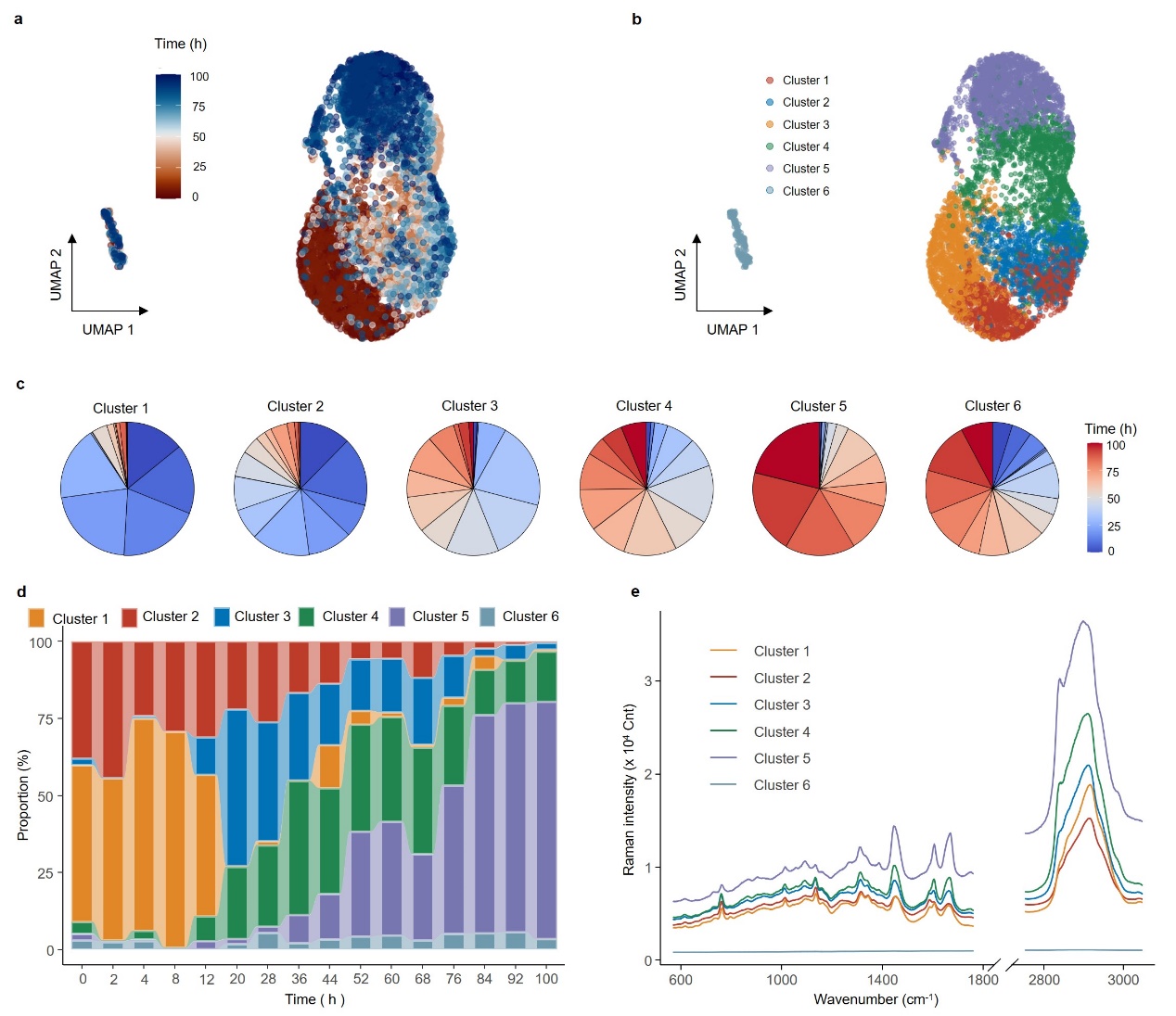

**Supplementary figure 8**. **“Phenotype analysis” module for the data matrix managed by SNR**. **a**, UMAP visualization of the data matrix managed by SNR. **b**, Louvain clustering results. **c-d**, Time point composition of each cluster and cluster composition at each time point. Cluster 6 appears evenly distributed across all time points. **e**, Mean spectra of four clusters, with cluster 6 consisting of spectra from null acquisitions.

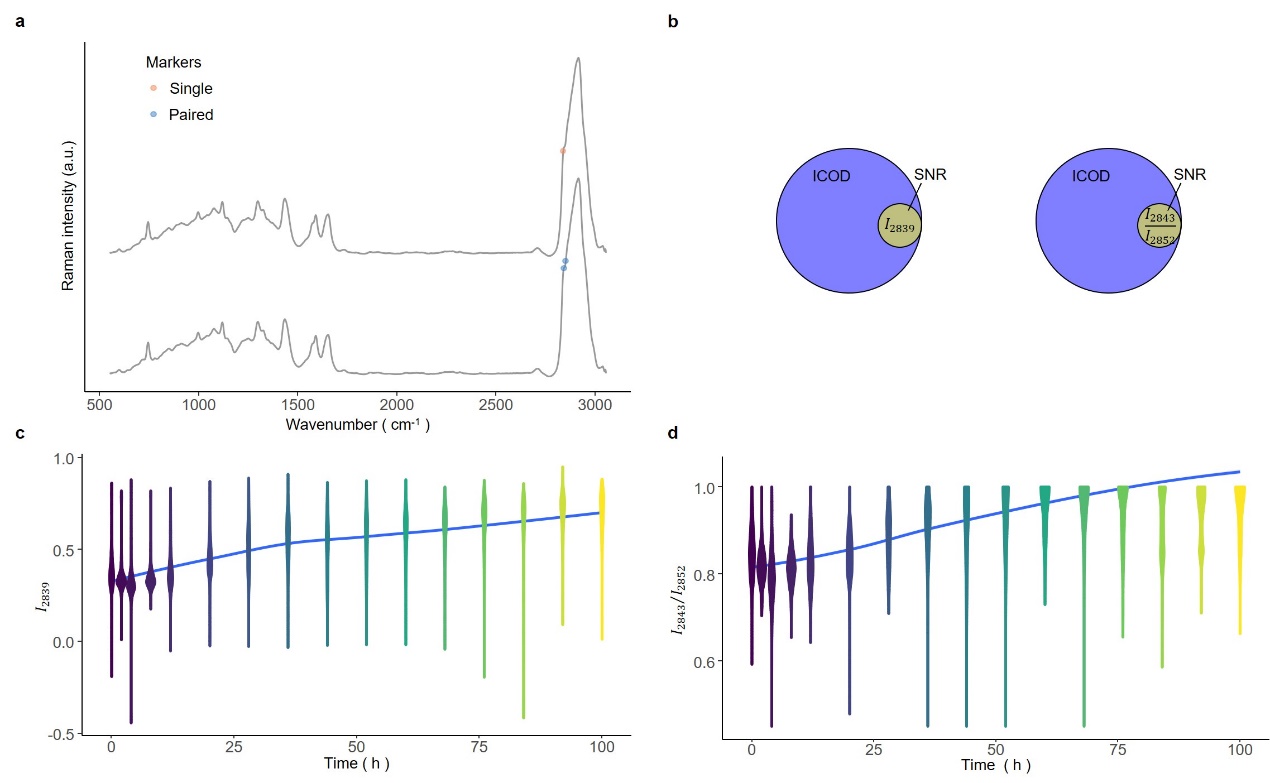

**Supplementary figure 9**. **“Raman markers” module for the data matrix managed by SNR**. **a**, Selected markers identified using the “correlation” method. **b**, Fewer markers were found compared to the ICOD-managed dataset. **c-d**, Intensity changes of singular and paired markers over time, showing more overlap compared to markers identified in the ICOD-managed dataset.

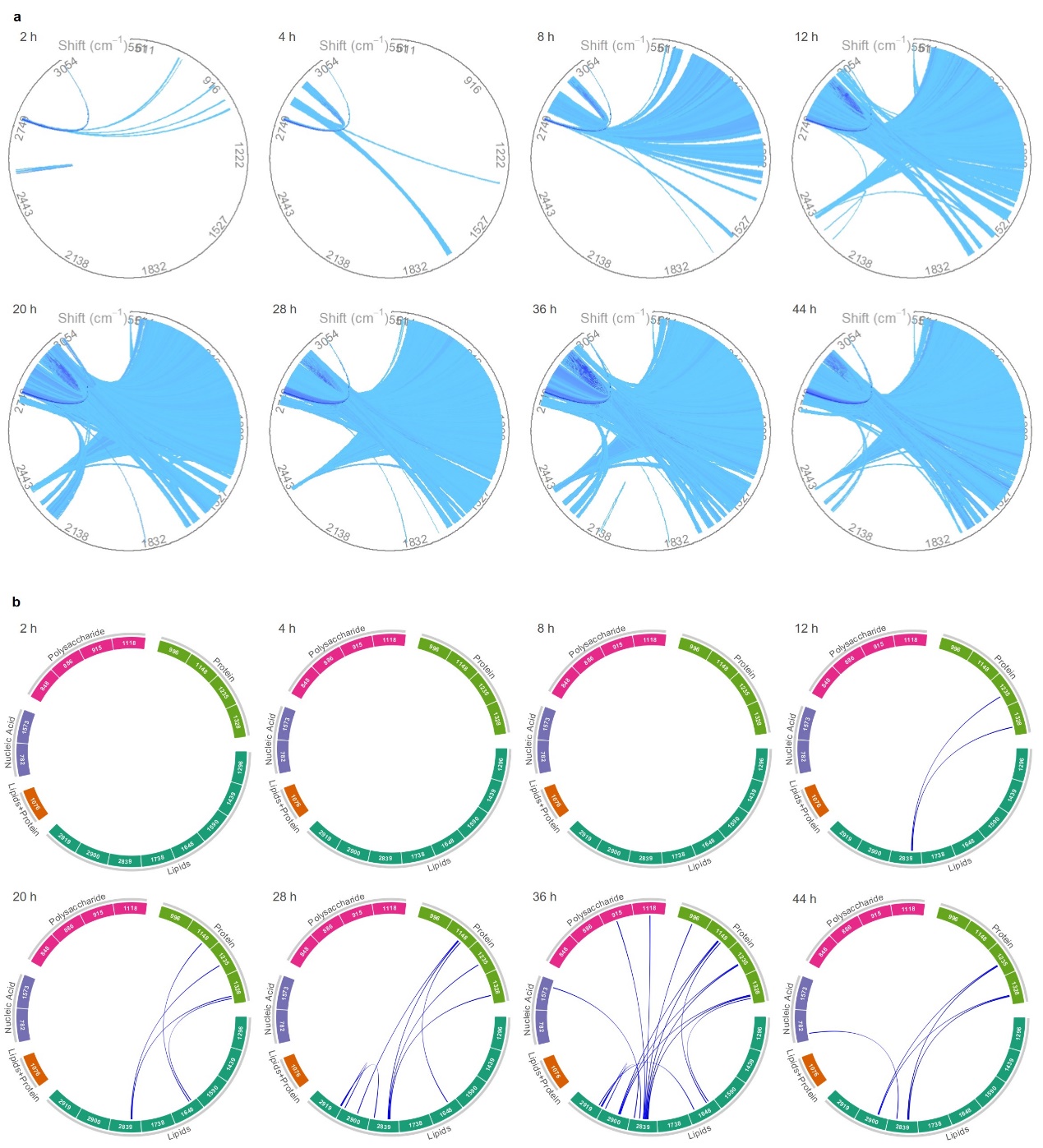

**Supplementary figure 10**. **“Intra-Ramanome” module for the data matrix managed by SNR**. **a**, Global-IRCN. **b**, Local-IRCN. Connections between Raman bands are colored based on their correlations. These IRCNs showed some confusion in metabolite conversions.

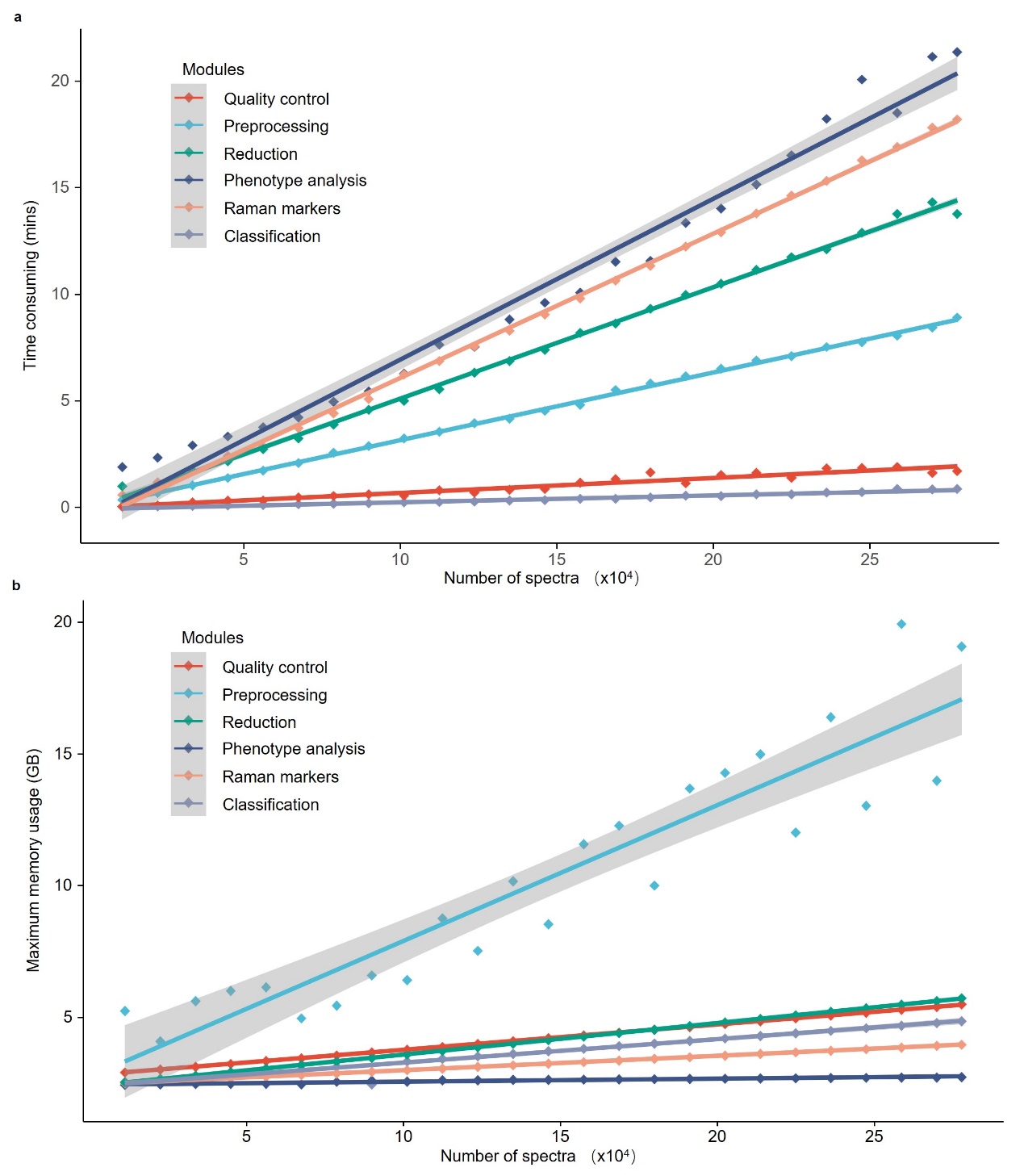
**Supplementary figure 11**. **Runtime and maximum memory usage with increasing data volume for each module in RamEx**. **a**, Run time of each module as data volume increases. **b**, Maximum memory usage of each module as data volume increases.

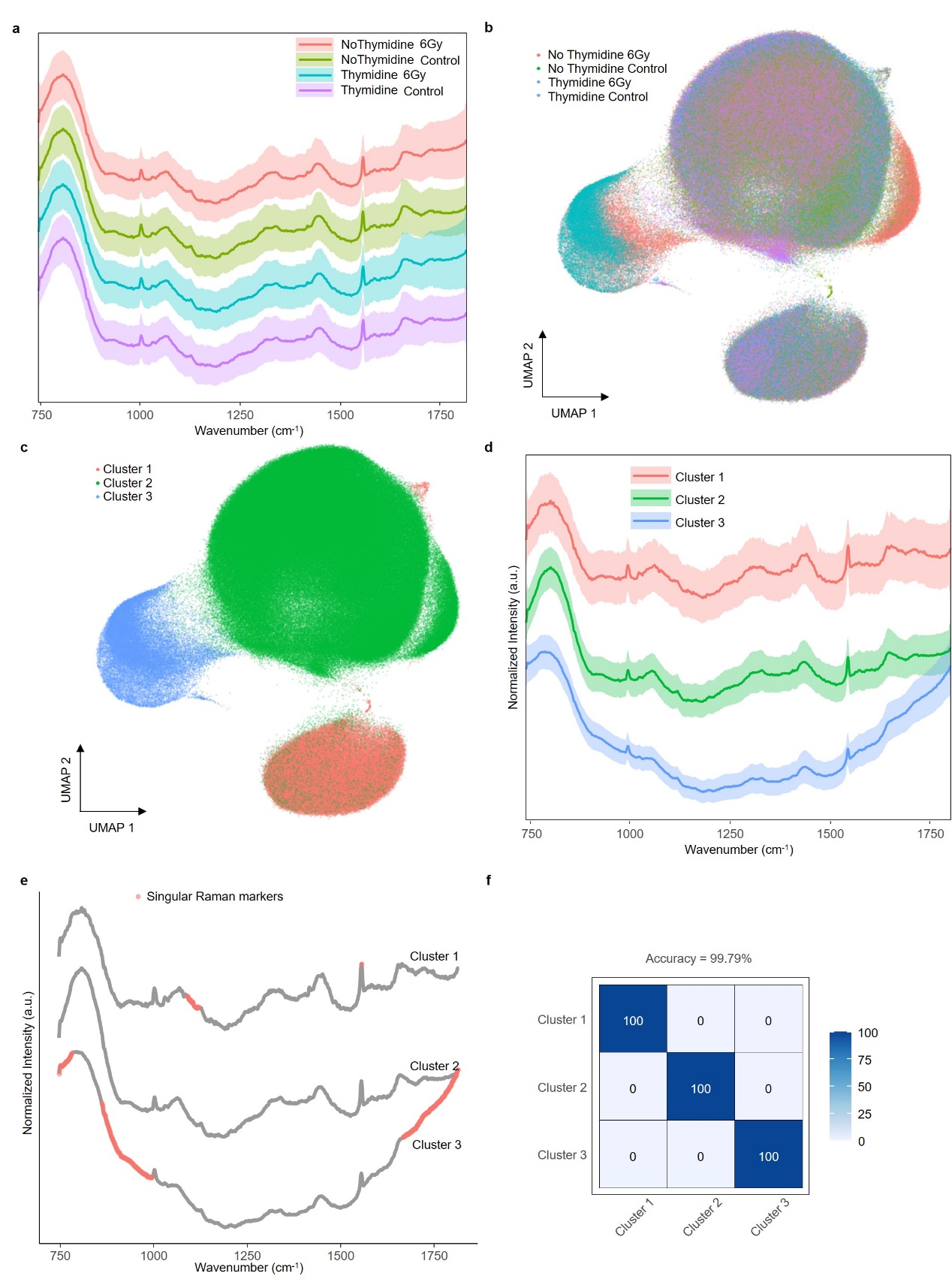
**Supplementary figure 12**. **Performance of RamEx on a dataset of about 800,000 spectra**. **a**, Mean spectra of the data matrix after preprocessing and ICOD’s quality control, points are colored by given labels. **b**, “Feature reduction” module. **c**, “Phenotype analysis” module. **d**, Mean spectra of the identified phenotypes. **e**, “Raman markers” module. **f**, “Classification” module.

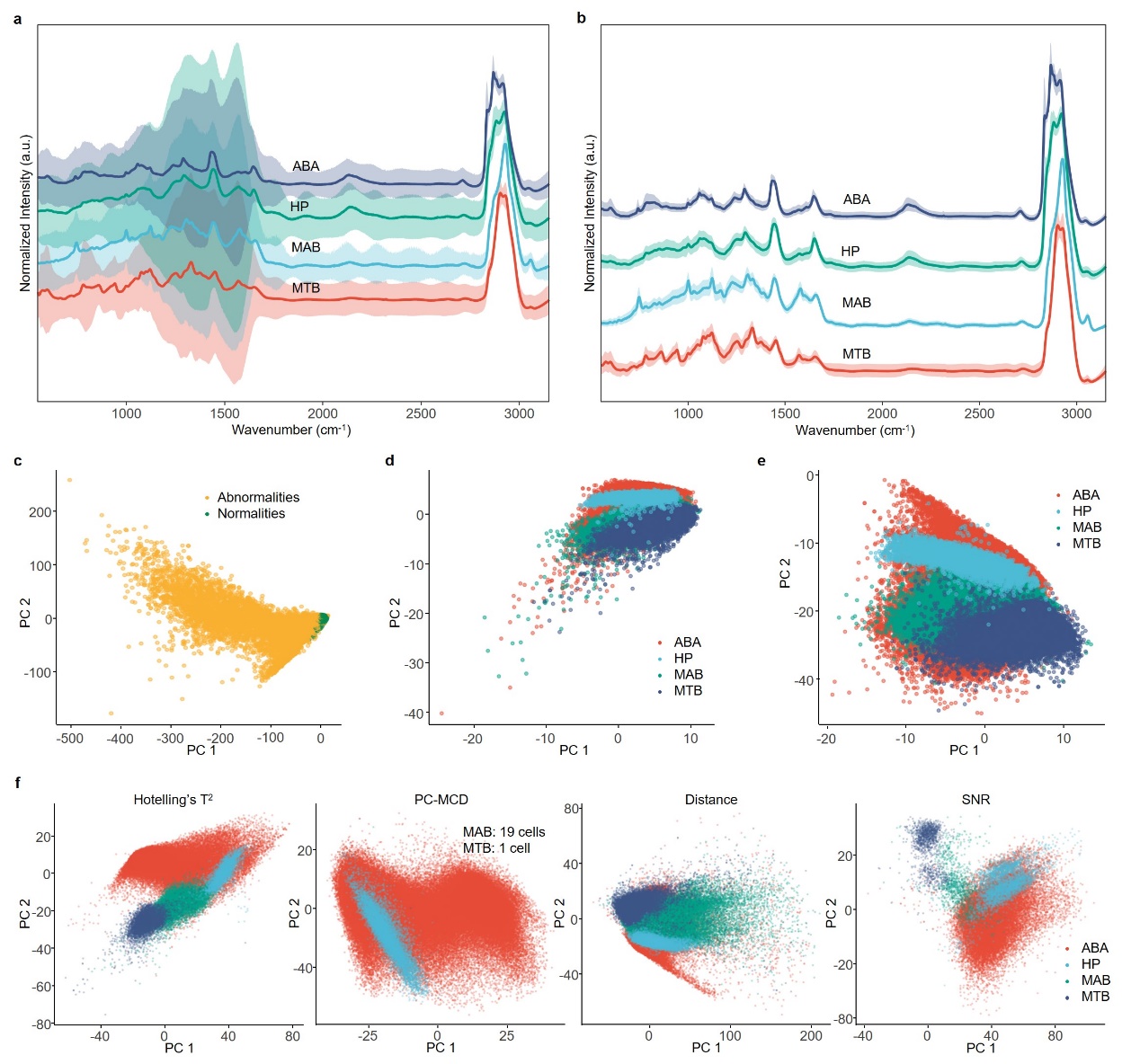
**Supplementary figure 13**. **PCA visualizations of the 4-species dataset managed by five different methods**. **a**, Mean spectra and standard deviations of the uncontrolled dataset, exhibiting large intensity variance among cells. **b**, Mean spectra of the dataset managed by the ICOD method. **c**, PCA visualization of the uncontrolled dataset, colored by quality as defined by ICOD, with most of the variance in the matrix originating from outliers. **d**, Valid spectra projected into the principal component space shown in **c**. **e**, PCA visualization of the ICOD-managed dataset. **f**, PCA reductions of datasets managed by other methods: Hotelling’s T^2^, PC-MCD, Euclidean distance, and SNR, respectively.

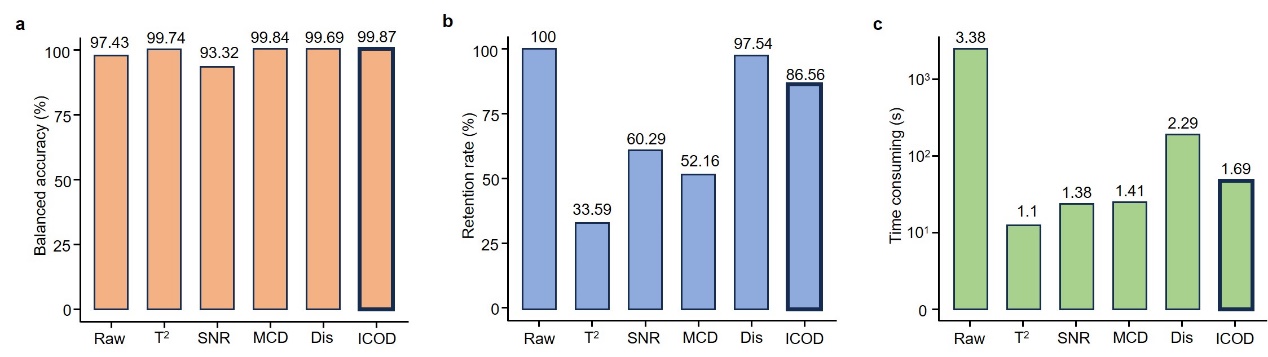
**Supplementary figure 14**. **Performance of the “Classification” module for datasets managed by different methods**. **a**, Balanced accuracy of validation sets, with single-cell spectra randomly divided into a training set and a test set at 7:3 ratio. ICOD achieved the highest prediction accuracy. **b**, Proportion of residual spectra retained by different methods. Overly strict methods like Hotelling’s T^2^, PC-MCD, and SNR retain only about 60% of the data, leading to significant resource wastage. **c**, Time required for SVM model learning on train sets. Distance-based method required longer model training times, suggesting slower SVM convergence and difficulty in identifying true features.

1.	B, H., ChemoSpec: Exploratory Chemometrics for Spectroscopy 2024.

2.	Beleites, C., A. Bonifacio, M. Dahms, B. Egert, S. Fuller, V. Gegzna, R. Guliev, B.A. Hanson, M. Hermes, M. Kammer, R. Kiselev, S. Mellor, E. Oduniyi, and V. Sergo, hyperSpec: a package to handle hyperspectral data sets in R. 2024.

18.	Guo, S., J. Popp, and T. Bocklitz, Chemometric analysis in Raman spectroscopy from experimental design to machine learning-based modeling. *Nat Protoc*, 2021.

32.	Etemadi, S. and M. Khashei, Etemadi multiple linear regression. *Measurement*, 2021. 186.

33.	Nelder, J.A. and R.W.M. Wedderburn, Generalized Linear Models. *Journal of the Royal Statistical Society. Series A (General)*, 1972. 135(3).

34.	Teng, L., X. Wang, X.J. Wang, H.L. Gou, L.H. Ren, T.T. Wang, Y. Wang, Y.T. Ji, W.E. Huang, and J. Xu, Label-free, rapid and quantitative phenotyping of stress response in E. coli via ramanome. *Scientific Reports*, 2016. 6.

35.	Blondel, V.D., J.-L. Guillaume, R. Lambiotte, and E. Lefebvre, Fast unfolding of communities in large networks. *Journal of Statistical Mechanics: Theory and Experiment*, 2008. 2008(10).

36.	MacQueen, J. *Some methods for classification and analysis of multivariate observations*. in *Proceedings of the Fifth Berkeley Symposium on Mathematical Statistics and Probability*. 1967.
